# lisaR: An LLM-Inferred Semantic Annotation of biological categories for gene set enrichment analysis

**DOI:** 10.64898/2026.09.29.755425

**Authors:** Claudia Velasco-López, Guillermo de la Vega-Barranco, David Olmeda

**Affiliations:** Drivers and Biomarkers of Metastasis Laboratory, Cancer Program, Instituto de Investigaciones Biomédicas Sols-Morreale (IIBM), CSIC-UAM, Madrid, Spain; Melanoma Group, Spanish National Cancer Research Centre (CNIO), Madrid, Spain

## Abstract

Gene set enrichment analysis (GSEA) turns differential expression results into lists of enriched gene sets. These lists are often long and redundant and span several gene-set collections, which makes their biological interpretation difficult. We present lisaR, an R package that reads previously computed differential expression or differential abundance results, runs GSEA and organises the output with the LISA dictionaries (LLM-Inferred Semantic Annotation). The dictionaries place 13,511 gene sets from the three Gene Ontology branches and from curated pathway collections such as Reactome, WikiPathways, KEGG, BioCarta and PID in 158 biological categories grouped into 42 supercategories within four dictionaries. Four open-weight language models scored the semantic fit of each gene set to the categories, and a consensus rule selected the assignments. The dictionaries are distributed with the package and do not depend on the data: studies analysed at different times with the same dictionaries are interpreted with the same categories, and no language model is queried during an analysis. For each category, lisaR summarises the direction of its significant gene sets and applies closed testing with Hommel’s robust procedure, which states whether the category shows evidence of enrichment and gives a simultaneous lower bound on how many of its gene sets are enriched. lisaR results are summarised in a report that opens in a web browser and leads from each category to its gene sets and genes, with every figure saved together with its data and R code. We illustrate the workflow with RNA-seq from melanoma biopsies taken before and during nivolumab treatment and with paired tumour and adjacent-tissue proteomics from clear cell renal cell carcinoma. Availability: lisaR is available at https://github.com/DBM-OlmedaLab/lisaR and archived at https://doi.org/10.5281/zenodo.23044930; documentation at https://olmedalab.org/lisaR/.

## Introduction

Gene set enrichment analysis relates the ranked results of a differential expression analysis to predefined groups of genes [1]. Gene Ontology and the curated pathway collections of MSigDB describe biological function at many levels of detail [2, 3], so a single comparison can return hundreds of significant gene sets that share genes and overlap in meaning. Deciding which processes such a list points to, and how strongly, is left to the researcher, and it often takes hours of careful analysis.

Several tools help with this step, and they follow a few recognisable strategies (Table 1). The first reduces redundancy among related terms. REVIGO keeps a representative of each group of semantically similar GO terms, and orsum discards a term when a more significant term annotates at least the same genes, which also lets it filter several enrichment results together [4, 5]. clusterProfiler offers a semantic simplification of GO results, and simplifyEnrichment clusters functional terms and summarises each cluster with word clouds [6, 7]. A second strategy organises gene sets into networks. EnrichmentMap draws enriched gene sets as a network whose edges represent shared genes, vissE groups related gene sets and displays the molecular data behind them, and GeneFEAST builds communities of terms around the genes of interest [8, 9, 10]; GeneTonic combines such networks with gene-level measurements in an interactive application [11]. A third strategy organises results around biological themes fixed before the analysis. GO subsets, or slims, are reduced sets of broad GO terms, Categorizer assigns genes to user-defined biological groups and tests the enrichment of each group, and thematicGO maps GO terms to themes through curated keywords [12, 13, 14]. A fourth looks across several analyses at once: mitch tests enrichment across many contrasts together, and simplifyEnrichment, GeneFEAST and SummArIzeR display the results of several gene lists or experiments side by side [15, 7, 10, 16].

**Table 1.**
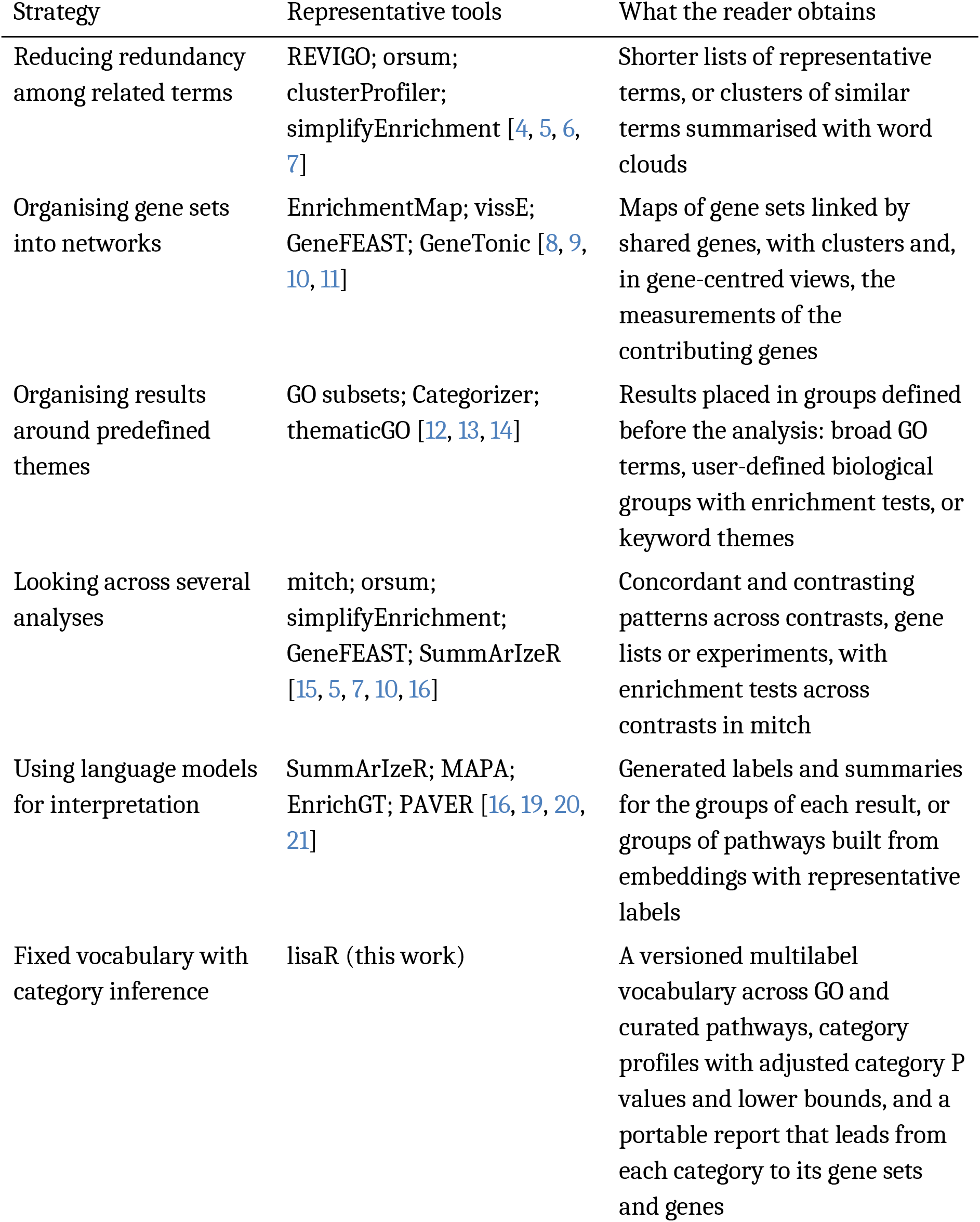
Strategies for organising and inspecting enrichment results. A tool can follow several strategies. Supplementary Section S7 describes ten of these tools from their publications.

Large language models open a fifth route. Hu and colleagues found that GPT-4 proposed functions similar to the curated name for 73% of GO gene sets, whereas the other models they tested gave confident answers for random gene sets [17]. GeneAgent reduces unsupported statements by checking its own output against biological databases [18]. For enrichment results, SummArIzeR, MAPA and EnrichGT use language models to annotate the groups of terms or pathways found in each analysis, and PAVER groups pathways by precomputed embeddings and labels each group with a representative pathway [16, 19, 20, 21].

These tools have made enrichment results much easier to read, and each answers a different need. Their designs also shape what a summary keeps. Reduction keeps one representative term for each group of similar terms, and the reader returns to the full list for the rest. The theme vocabularies fixed in advance that we cite are built on Gene Ontology, so the curated pathway collections fall outside them. Groups built from each result adapt to that result, so the same process can be grouped differently in two studies, and following a pattern across studies is simpler when every study is interpreted with the same categories. lisaR uses language models at a different point: once, to build a vocabulary that is then kept fixed. We defined biological categories and their scopes, and scored against them the gene sets of the three Gene Ontology branches (biological process, molecular function and cellular component) and of the canonical pathway collections of MSigDB, which include Reactome, WikiPathways, KEGG, BioCarta and PID [2, 3, 22, 23, 24]. Four language models provided the scores, and a consensus rule selected the assignments. The released LISA dictionaries place 13,511 gene sets in 158 categories grouped into 42 supercategories, with one dictionary for each of four collections: GOBP-C2, GOMF, GOCC and PATHWAYS. Because the categories do not depend on the data, studies analysed at different times with the same dictionaries are interpreted with the same categories, and no language model is queried when a study is analysed. On this vocabulary, lisaR adds a descriptive profile of the significant member sets of each category and category-level inference by closed testing, which states whether a category shows evidence of enrichment and gives a simultaneous lower bound on how many of its member sets are enriched [25]. Its portable report leads from each category to its gene sets and genes. Below we describe the package and its methods, and apply them to a melanoma RNA-seq cohort and to a renal cancer proteomic cohort.

## Methods

### Workflow and inputs

lisaR is an R package for R 4.1 or later, distributed under GPL-3.0-or-later. Every study follows the same path (Fig. 1B). The input is the complete table of a previously fitted differential expression or differential abundance analysis for one declared comparison, with gene identifiers, effect sizes, a signed ranking statistic, and gene-level P values and adjusted P values. lisaR reads this table as a TSV or CSV file or as an R data frame, for example exported from limma, and it reads DESeq2 results and edgeR result objects directly [26, 27]. A DESeq2 Wald statistic and a moderated limma t statistic are typical rankings, and an expression or abundance matrix can be added for gene-level displays. Gene-set memberships are registered once per species and identifier system. For human data, the function prepare_lisa_msigdb_resource() builds them from MSigDB release 2026.1, and memberships for mouse or other systems can be registered with install_lisa_resource() [28].

**Figure 1.**
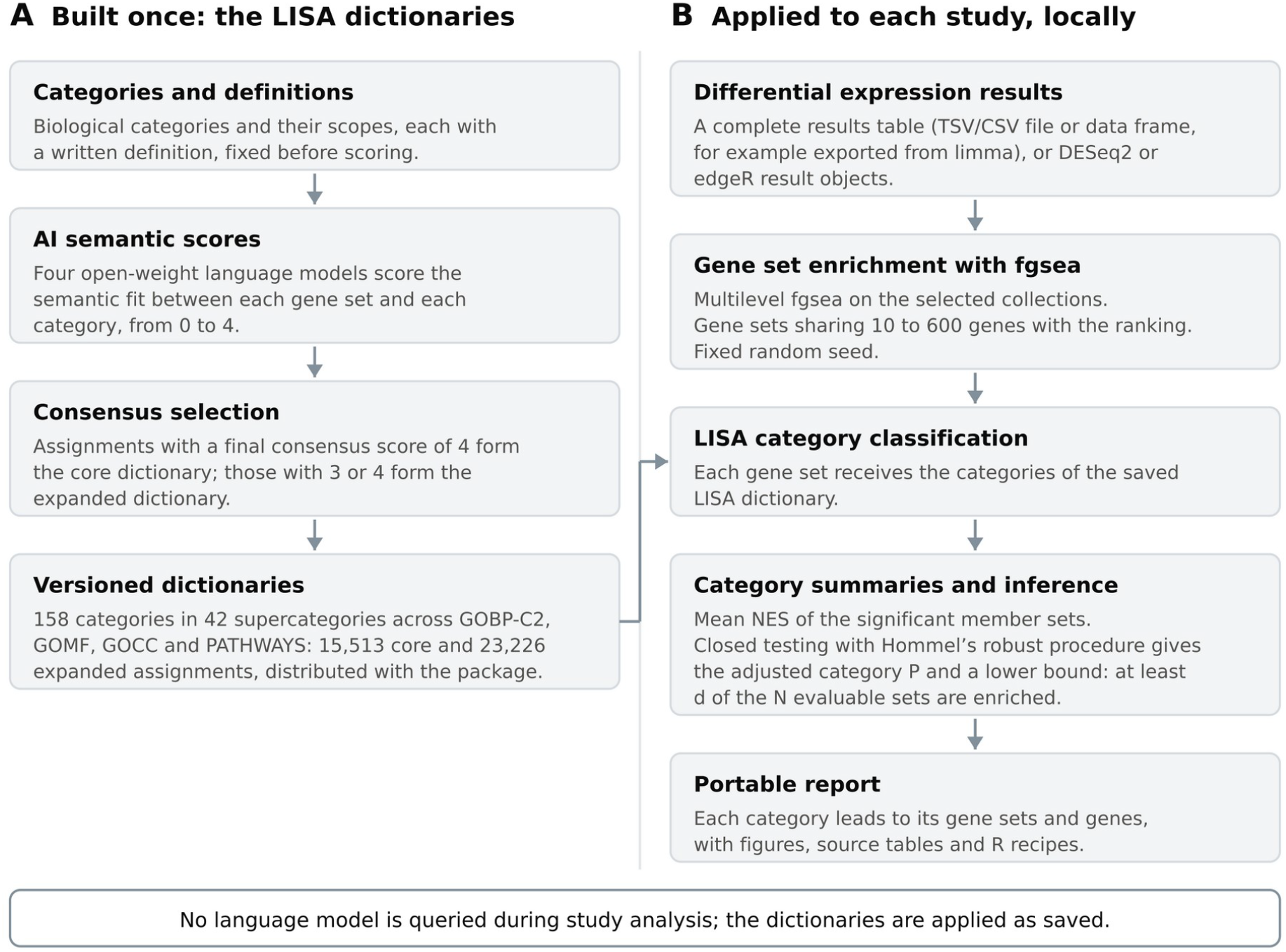
The lisaR workflow. **A**. The LISA dictionaries are built once. The categories and their definitions are fixed first; four open-weight language models score the semantic fit of each gene set to the categories; a consensus rule selects the assignments, with a final consensus score of 4 for the core dictionary and of 3 or 4 for the expanded dictionary; and the dictionaries are distributed with the package. **B**. Each study is analysed on the local computer. lisaR reads complete differential expression results, runs GSEA with fgsea, classifies the gene sets with the saved LISA dictionaries, summarises and tests each category, and writes a portable report. No language model is queried during the analysis of a study.

The function run_lisa() then ranks the genes, runs GSEA on the selected collections, classifies the gene sets with the LISA dictionaries, computes the category summaries and the category inference, and writes the report. Table 2 lists the main functions.

**Table 2.**
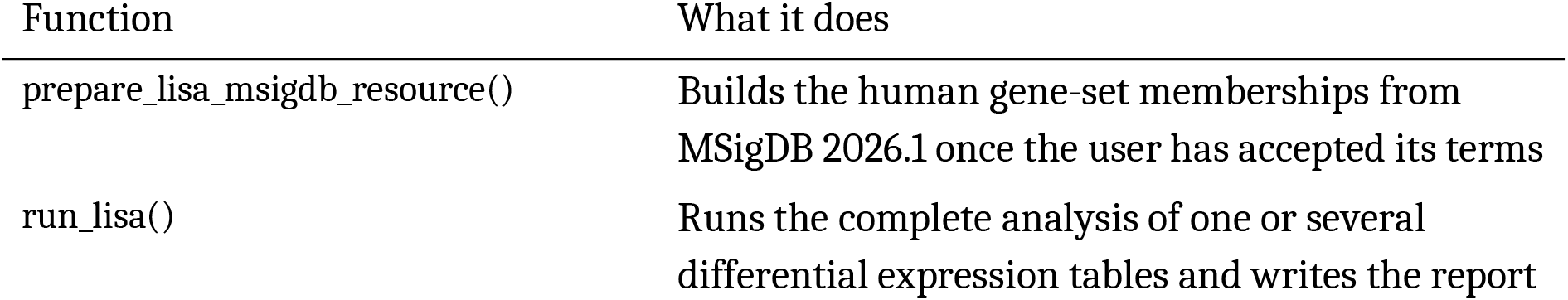
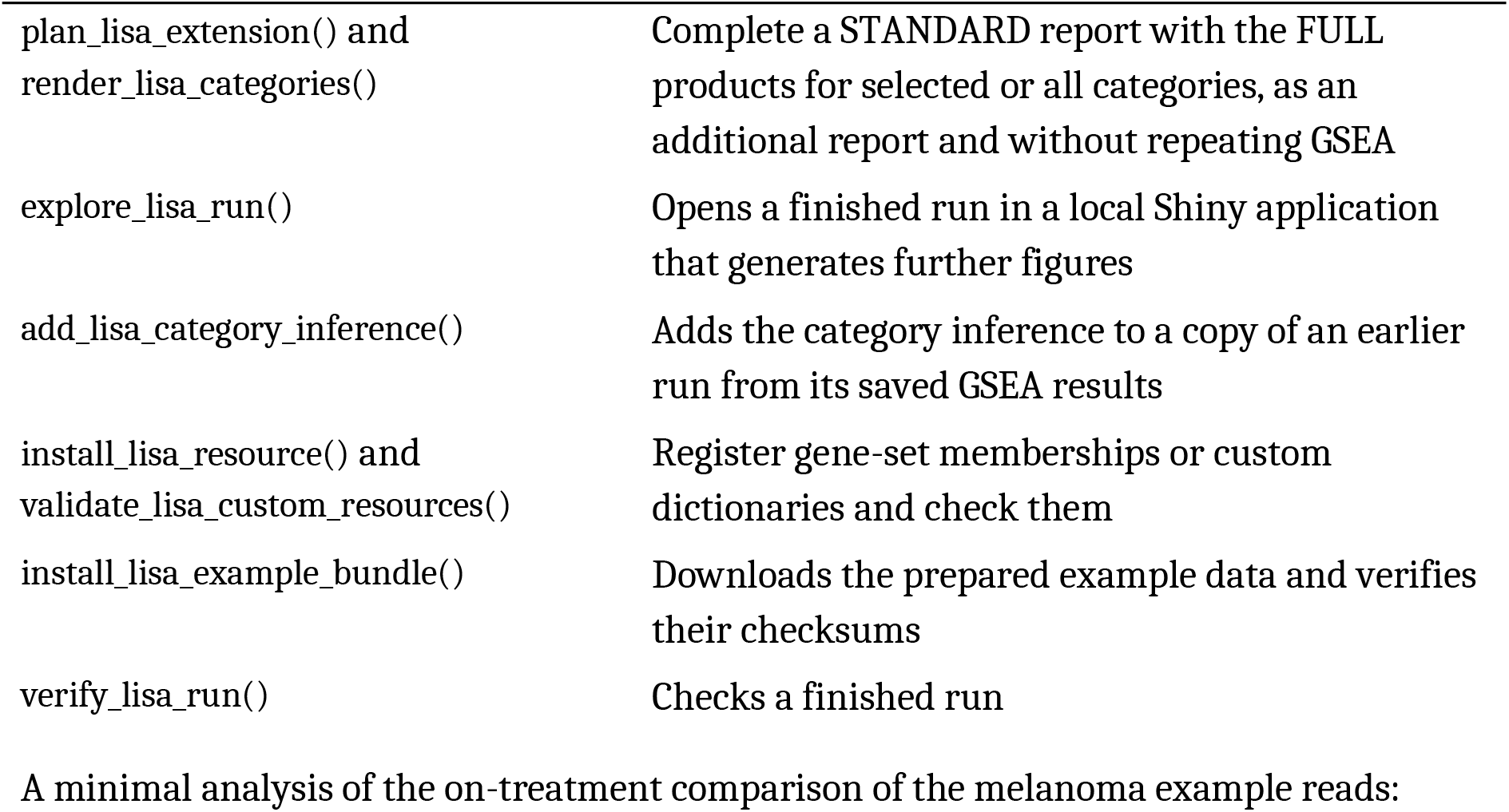
Main functions of lisaR.

A minimal analysis of the on-treatment comparison of the melanoma example reads:

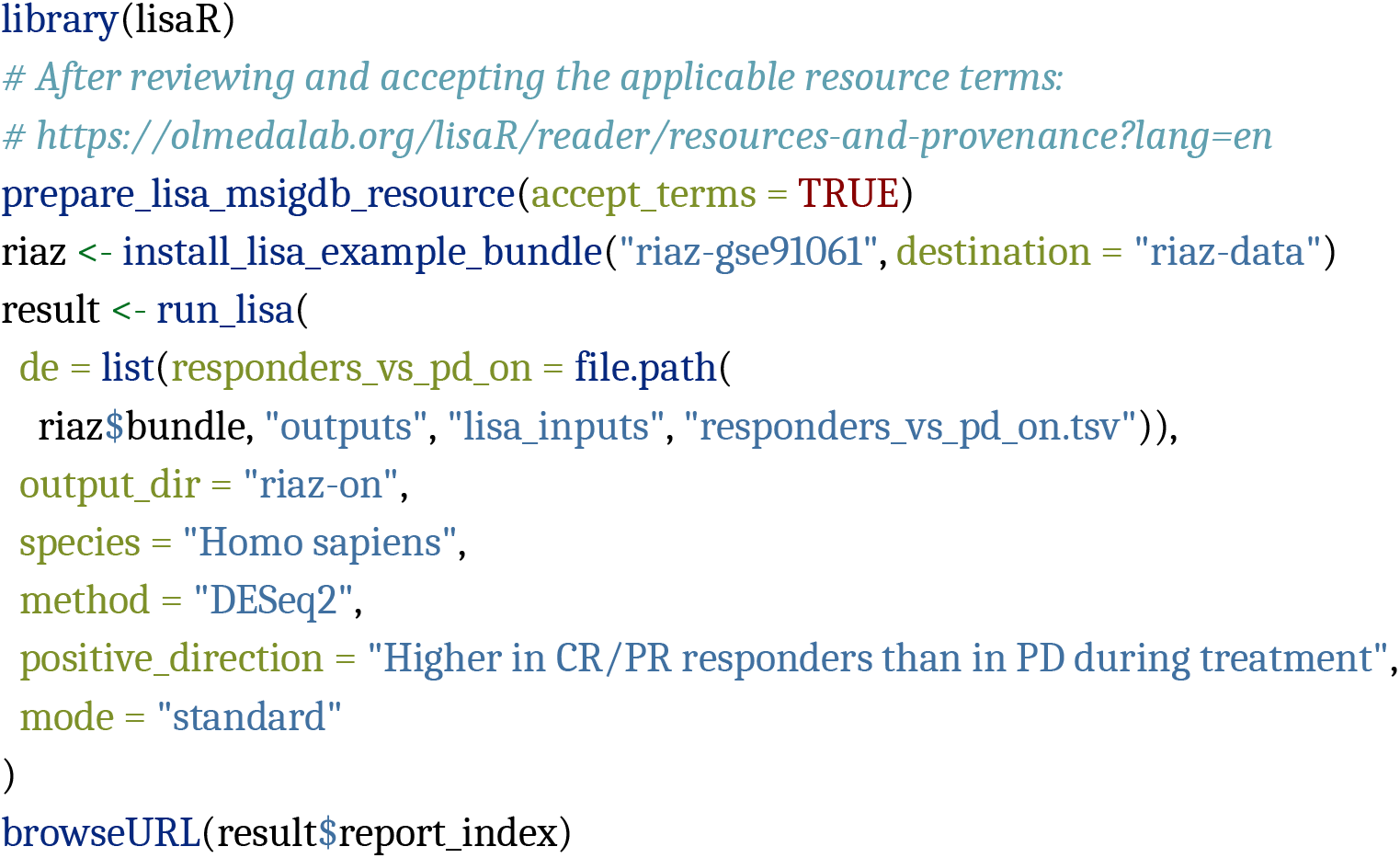

### The LISA dictionaries

#### Collections and categories

The dictionaries cover four collections. GOBP-C2 combines the Gene Ontology Biological Process gene sets with the canonical pathway gene sets of MSigDB C2 (C2:CP), from BioCarta, KEGG, PID, Reactome, WikiPathways and three smaller sources [2, 3, 24, 22, 23]. GOMF describes molecular activities, and GOCC cellular locations, structures and complexes. PATHWAYS reads the gene sets of GOBP-C2 through a narrower lens: it classifies only the sets whose names identify one of 69 named signalling systems. Hallmark gene sets can be analysed directly, without a LISA dictionary [29]. We defined the biological categories and their scopes, each with a written definition, before any scoring. The released dictionaries contain 158 categories grouped into 42 supercategories for display, and a gene set can belong to more than one category (Supplementary Section S1).

#### Construction

Four open-weight language models, Qwen3.5 27B, Gemma 4 31B, Mistral Small 3.2 24B and Phi-4 14B, scored the semantic fit between the gene sets of each collection and its categories on an ordinal scale from 0 to 4 (Supplementary Section S2). For GOMF, 0 denotes no relationship, 3 a major semantic relation and 4 a primary or canonical one; GOCC uses 0, 3 or 4 to judge compartment identity. A consensus rule combined the four scores into a final consensus score for each gene set and category. A final score of 4 places the assignment in the core dictionary, and a score of 3 or 4 in the expanded dictionary, which therefore contains the core. PATHWAYS adds a name-first condition: the gene-set name and its source must identify the signalling system, and canonical genes can only reinforce that reading. Supplementary Section S2 gives the rule in full and the treatment of missing scores.

#### Evaluation

We evaluated the dictionaries on 217 gene-set/category relations. The set contains 198 core assignments of 113 gene sets, which together cover 145 of the 158 categories, and 19 negative controls that pair gene sets without an assignment with categories chosen independently of their content. It was drawn from the human MSigDB gene sets other than Hallmarks, after excluding the 1,388 gene sets used while developing the construction rules, and its rows were presented in random order (Supplementary Section S3). Three evaluators, working independently, judged every relation as central, secondary, unsupported or indeterminate. The three evaluators supported all 198 assignments and rejected all 19 controls.

Four further language models, none of which took part in the construction, judged the same relations: Claude Sonnet 5 and Claude Opus 5 (Anthropic) and GPT-5.6 Terra and GPT-5.6 Sol (OpenAI), with two runs per model. A relation counted as rejected when at least four of its eight judgements were unsupported. The models supported 196 of the 198 assignments in all eight judgements and rejected all 19 controls. They disagreed on two assignments.

GOMF_NUCLEOSOMAL_DNA_BINDING in Structural constituent activity received seven supporting judgements of eight. GOMF_DENEDDYLASE_ACTIVITY in Ubiquitin and SUMO ligase activity received two and was rejected: the models that rejected it distinguished the removal of NEDD8 from the ligase activity named by the category.

### Enrichment and category analysis

#### Gene set enrichment

lisaR ranks all the genes of the input table by their signed statistic, so that genes that are not significant on their own still contribute to the enrichment. It then runs GSEA with fgsea on each selected collection, using the gene sets that share between 10 and 600 genes with the ranking, the default limits used in the examples [1, 30]. All results are kept, including those of the gene sets without a LISA category. Supplementary Section S5 describes how identifiers are matched and how duplicated genes are handled. An over-representation analysis of a thresholded gene list against a declared background is available as a separate, optional route.

#### Category summaries

For each category, lisaR summarises the member sets that have a finite NES and a GSEA adjusted P value at or below a configurable cutoff, 0.25 in the examples. The mean NES is the arithmetic mean over these sets; each set counts once per category, and a set assigned to several categories contributes to each of them. The summary reports n, the number of significant member sets, and how many of them have positive or negative NES. The mean NES and its sign counts describe the direction and size of the significant results. Whether the category as a whole shows evidence of enrichment is the question answered by the inference.

#### Category inference

Category inference applies closed testing with the robust variant of Hommel’s procedure, which controls the familywise error rate under any dependence between gene sets [31, 32, 25], as implemented in the hommel R package [33]. It uses the raw GSEA P values of the member sets, independently of the 0.25 cutoff and of the direction of the NES. For each analysis, the hypothesis family contains every eligible gene set across the four collections, including the sets without a LISA category and excluding Hallmarks. For a category with N evaluable member sets, the procedure gives an adjusted category P value, and a significant category contains at least one enriched member set. At the 5% level, the same procedure gives d, a simultaneous 95% lower bound on the number of enriched member sets: at least d of the N sets, a proportion of at least d/N, are enriched. Because the bounds are simultaneous, they hold together for all the categories of an analysis and can be read side by side. Figures mark the adjusted category P with the usual thresholds: **** ≤ 0.0001, *** ≤ 0.001, ** ≤ 0.01 and * ≤ 0.05. Supplementary Section S5 gives the details, with a worked example.

#### Comparison of two profiles

lisaR can place two completed category profiles side by side, A and B, and report the difference between their category means, A minus B. Each side keeps its own significant sets. When only one side has significant sets, the other can be shown as a labelled contextual value computed from its evaluable sets (Supplementary Section S5).

### The report and custom dictionaries

#### The report

The report is a set of linked web pages that can be opened in a browser and shared as it is, without R, Shiny or a network connection. For each analysis and collection, it shows the category profile together with the category inference. Each category opens on its member gene sets, with their NES, adjusted P values and leading edges, and on the genes behind them, with their effects and adjusted P values. Every figure is saved as PNG and SVG with its source table and an R recipe that redraws it.

The two report modes use the same statistics and differ in what they draw (Table 3). STANDARD, the default, writes the complete tables, the category profiles and inference, an evidence page for every category and, when two profiles are compared, their aligned category, gene-set and gene evidence. FULL adds detailed displays of the gene sets and genes of every category with at least one significant member set. A STANDARD report can be completed later without repeating GSEA. plan_lisa_extension() and render_lisa_categories() write the FULL products for selected or all categories into an additional report, and explore_lisa_run() opens the run in a local Shiny application that generates further figures one by one and exports them with their data and recipes. Both routes leave the original run unchanged.

**Table 3.**
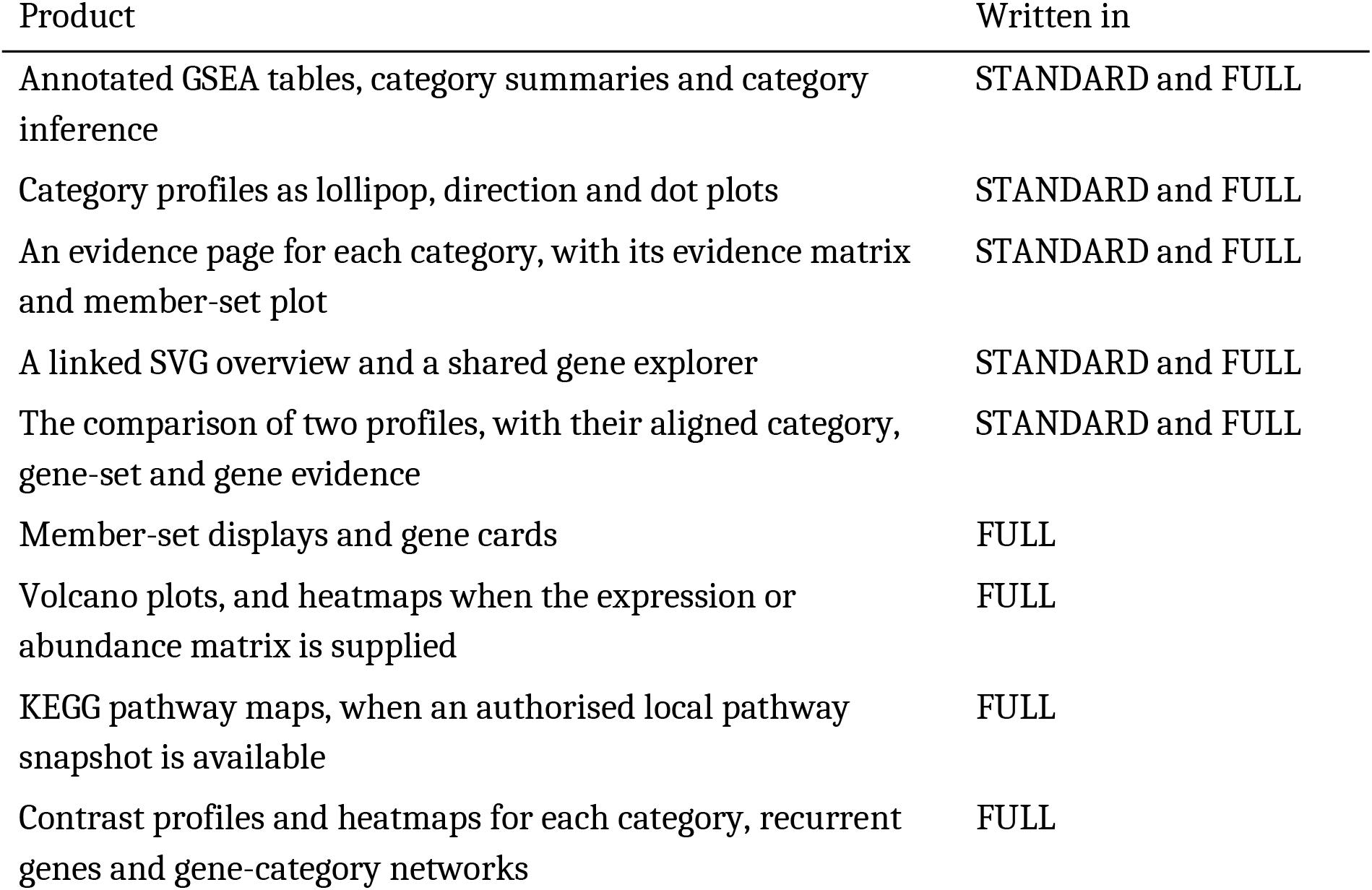
Products of the two report modes.

#### Gene-level evidence

The report links genes to gene sets through the GSEA leading edge, the genes that drive the enrichment score of a set, and through complete memberships when these are supplied. A gene card ranks the genes associated with a category by two descriptive scores. LISA support combines the number of classified sets whose leading edge contains the gene with the absolute value of their mean NES and their minimum adjusted P value; the DE-weighted contribution combines LISA support with the absolute log2 fold change of the gene and its DE adjusted P value. Both scores are prioritisation heuristics, defined in Supplementary Section S5.

#### Custom dictionaries

Users can apply their own categories in the same workflow. A custom dictionary consists of three tables: the gene-set memberships, the assignments of gene sets to categories, and a category map that groups the categories into supercategories and sets their order. install_lisa_resource() registers the tables with their checksums, validate_lisa_custom_resources() checks that every assignment refers to a known category and to a gene set present in the memberships, and run_lisa() uses them through its resources argument. Supplementary Section S4 gives the required columns and the validation rules, and the package vignette “Custom dictionaries and supercategories” works through an example step by step.

### Example studies

We chose two public studies that differ in molecular layer and design. The RNA-seq example uses the melanoma cohort of Riaz et al. (Cell, 2017), treated with nivolumab (GEO accession GSE91061) [34]. Its biopsies were taken before and during treatment in patients with a recorded clinical response, which allows responders and non-responders to be compared at two time points. We kept the patients with one pretreatment and one on-treatment biopsy, a complete or partial response (CR/PR) or progressive disease (PD), and a known history of prior ipilimumab, and excluded the published PCA outlier Pt3. This left 26 patients and 52 biopsies, eight with CR/PR and 18 with PD. Separate DESeq2 models for each time point compared CR/PR with PD, adjusting for prior ipilimumab, and their Wald statistics ranked the genes [26]. The same upstream analysis also fitted a patient-blocked longitudinal model with a formal time-by-response interaction (Supplementary Section S6). The on-treatment GOBP-C2 input had 22,070 rows and 21,520 ranked symbols. The report compares the on-treatment profile (A) with the pretreatment profile (B).

The proteomic example uses the clear cell renal cell carcinoma cohort of Clark et al. (Cell, 2019), from the Clinical Proteomic Tumor Analysis Consortium (CPTAC study PDC000127) [35]. It applies the same categories to another molecular layer, protein abundance, in a paired design with tumour and adjacent non-tumour tissue (NAT) from the same patients, and its input comes from limma rather than DESeq2. From the processed abundance table, we kept 80 patients with matched tumour and NAT, 160 samples in total. RefSeq features were mapped conservatively to genes, genes represented by several rows were excluded, and complete paired measurements were required, which left 6,482 genes. For each gene, the within-patient tumour-minus-NAT differences were fitted with an intercept-only limma model with empirical Bayes moderation, and the moderated t statistic ranked the genes [27] (Supplementary Section S6). The run used the four LISA collections and the core dictionary.

## Results

### Contents and coverage of the LISA dictionaries

The core dictionaries place 13,511 distinct gene sets in the LISA categories: 6,937 from GO Biological Process, 1,768 from GO Molecular Function, 990 from GO Cellular Component, 1,701 from Reactome, 830 from WikiPathways, 805 from KEGG, 273 from BioCarta, 188 from PID and 19 from other sources (Supplementary Section S1). They contain 15,513 gene-set/category assignments in the core tier and 23,226 in the expanded tier, which includes the core (Fig. 2A). GOBP-C2, the broadest collection, assigns 10,743 gene sets to 40 categories in its core tier. A gene set can carry several labels, and 1,826 core gene sets have more than one. BIOCARTA_41BB_PATHWAY, named after the costimulatory receptor 4-1BB (TNFRSF9), reached a final consensus score of 4 for Cytokine and inflammatory signaling and of 3 for Immune core, which places it in the core tier of the first category and in the expanded tier of the second; the expanded dictionary also assigns it to Core cell signaling (Fig. 2B). In PATHWAYS, WP_T_CELL_RECEPTOR_AND_COSTIMULATORY_SIGNALING carries two core labels, T-cell receptor signaling and CD28 costimulation signaling, one for each component of the named pathway (Fig. 2C).

**Figure 2.**
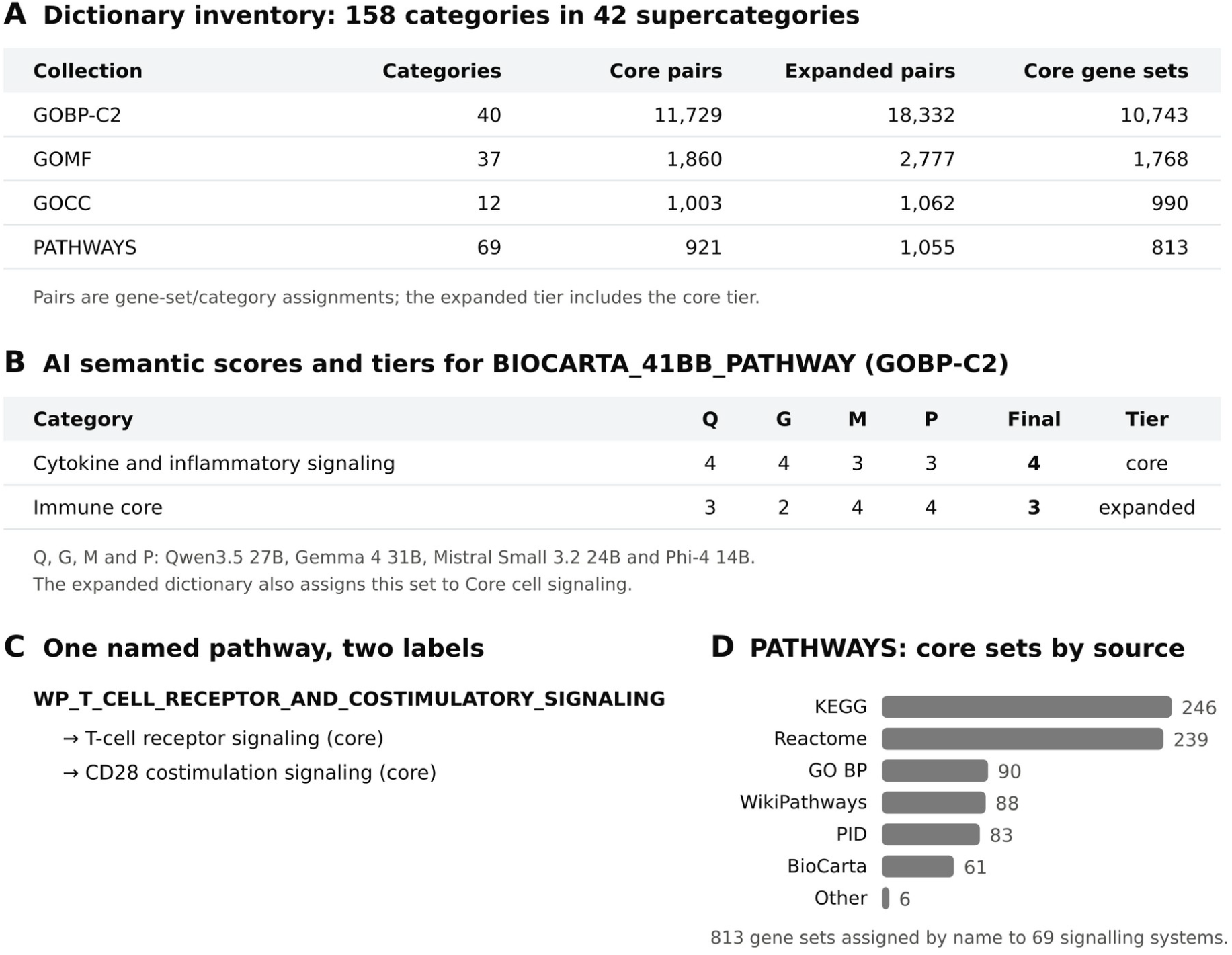
Contents of the LISA dictionaries. **A**. Inventory of the four collections. Pairs are gene-set/category assignments, and the expanded tier contains the core tier. Core gene sets are the distinct sets of each collection; the same set can appear in more than one collection. **B**. AI semantic scores of the four models for two categories of BIOCARTA_41BB_PATHWAY in GOBP-C2, with the final consensus score and the resulting tier. Q, Qwen3.5 27B; G, Gemma 4 31B; M, Mistral Small 3.2 24B; P, Phi-4 14B. **C**. A named pathway with two core labels in PATHWAYS. **D**. Sources of the 813 core PATHWAYS gene sets; “Other” groups the SA (3), SIG (2) and WNT (1) sets.

PATHWAYS is deliberately focused. Its core tier assigns 813 gene sets to 69 named signalling systems, mainly from KEGG (246), Reactome (239), GO Biological Process (90), WikiPathways (88), PID (83) and BioCarta (61) (Fig. 2D). Because it classifies a set only when its name identifies one of these systems, PATHWAYS gives a targeted reading of signalling alongside the broader GOBP-C2 categories.

In the three example analyses, the broad collections classified most of the gene sets that reached a GSEA adjusted P value of 0.25 or less. GOBP-C2 classified 93% of them: 3,857 of 4,142 in the on-treatment melanoma analysis, 2,485 of 2,676 before treatment and 2,576 of 2,759 in the renal cancer proteomics. GOMF classified 94 to 95% and GOCC 90 to 94%. The sets without a category remain in the complete tables and in the inference family.

### Category profiles of response in melanoma

The Riaz cohort comprises patients with advanced melanoma, some of whom had progressed on ipilimumab, who were biopsied within a week before their first dose of nivolumab and again, at the same site, between days 23 and 29 of the first cycle [34]. Of the 65 patients in the public data, 43 had a biopsy at both time points. Among them, we compared the two clearly separated outcomes of the best overall response by RECIST 1.1: objective response, complete or partial (CR/PR), and progressive disease (PD). Stable disease groups tumours that neither shrank enough to count as a response nor grew enough to count as progression, a range that includes both slow progression and lasting disease control, so the 15 patients with stable disease were not assigned to either group. One patient whose response could not be evaluated and the published PCA outlier Pt3 were also excluded, which left eight patients with CR/PR and 18 with PD. At each time point, DESeq2 compared CR/PR with PD, adjusting for prior ipilimumab, and lisaR analysed each comparison, with positive NES pointing towards CR/PR (Fig. 3A). The pretreatment comparison asks which programmes already distinguish the two groups before nivolumab, and the on-treatment comparison asks which distinguish them during treatment. Both profiles are interpreted with the same categories.

**Figure 3.**
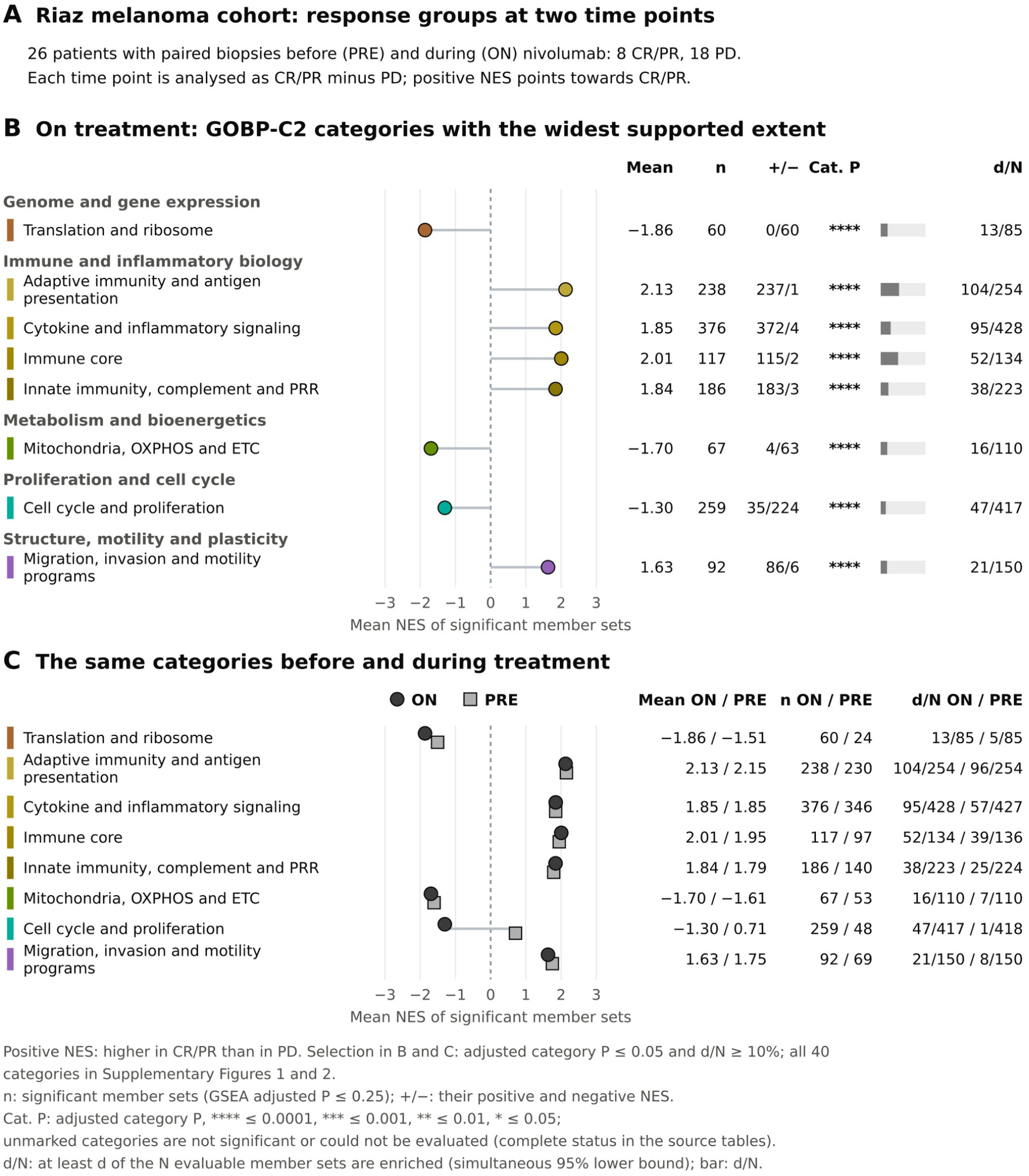
Category profiles of response to nivolumab in the Riaz melanoma cohort. **A**. Design: 26 patients with paired pretreatment (PRE) and on-treatment (ON) biopsies, eight with complete or partial response (CR/PR) and 18 with progressive disease (PD) [34]. Each time point is analysed as CR/PR minus PD, and positive NES points towards CR/PR. **B**. On treatment, the GOBP-C2 categories with an adjusted category P of 0.05 or less whose simultaneous lower bound covers at least 10% of their evaluable member sets, grouped by supercategory. Points show the mean NES of the significant member sets (GSEA adjusted P ≤ 0.25); n is their number and +/− counts their positive and negative NES. Cat. P gives the adjusted category P as stars. d/N states that at least d of the N evaluable member sets are enriched (simultaneous 95% lower bound), also drawn as a grey bar. **C**. The same categories on treatment (dark circles) and before treatment (light squares). The distance between the two points is a descriptive difference between the two profiles; the formal time-by-response interaction was estimated in the upstream model (Supplementary Section S6). All 40 categories are shown in Supplementary Figures 1 and 2.

On treatment, 30 of the 40 GOBP-C2 categories showed evidence of enrichment at an adjusted category P of 0.05 or less (Supplementary Figure 1). With so many categories passing this threshold, the lower bound is the more informative guide: it states how many member sets of each category are, at a minimum, enriched. Eight categories had lower bounds covering at least 10% of their evaluable member sets, and Figure 3B shows them.

As expected from earlier studies of this cohort and of PD-1 blockade in melanoma [34, 36], the four immune categories had the widest supported extent, all higher in CR/PR. For Immune core, at least 52 of its 134 evaluable member sets were enriched; its 117 significant member sets had a mean NES of 2.01, and 115 of them were positive, pointing towards CR/PR. Adaptive immunity and antigen presentation (at least 104 of 254 sets enriched), Cytokine and inflammatory signaling (95 of 428) and Innate immunity, complement and PRR (38 of 223) pointed in the same direction, with mean NES values between 1.84 and 2.13 and almost all significant sets positive. Migration, invasion and motility programs was also higher in CR/PR (at least 21 of 150; mean 1.63). Translation and ribosome (13 of 85), Mitochondria, OXPHOS and ETC (16 of 110) and Cell cycle and proliferation (47 of 417) pointed the other way: 60 of 60, 63 of 67 and 224 of 259 of their significant sets had negative NES, higher in PD.

PATHWAYS names the signalling systems within this immune signal. On treatment, 10 of its 69 categories showed evidence of enrichment, nine of them with every evaluable member set pointing towards CR/PR. T-cell receptor signaling had the widest supported extent, with at least 10 of its 16 evaluable sets enriched and all 16 with positive NES, followed by B-cell receptor signaling (at least 5 of 12) and IL12/IL23 signaling (2 of 8).

Because the categories are the same at both time points, the pretreatment profile can be read against the on-treatment one (Fig. 3C). Before treatment, 24 of the 40 categories showed evidence of enrichment (Supplementary Figure 2), and the immune categories already had similar profiles. Adaptive immunity and antigen presentation, for example, had a mean NES of 2.15 and a lower bound of 96 of 254 sets before treatment, against 2.13 and 104 of 254 during treatment, and the bound of Immune core went from 39 of 136 to 52 of 134. Cell cycle and proliferation differed between the two time points. Before treatment, its 48 significant sets had a mean NES of 0.71 and its lower bound was 1 of 418; on treatment, its 259 significant sets had a mean of −1.30 and the bound was 47 of 417. The comparison thus separates a persistent pattern, the immune signal of the responders, from a profile that changed, cell cycle and proliferation, which became higher in PD during treatment.

These patterns are consistent in direction with the original analysis of this cohort. Riaz et al. reported that the genes upregulated on therapy in responders covered a broad spectrum of immune functions, including lymphocyte activation, chemotaxis, cytokine signalling and cytolytic activity, and that the genes downregulated in responders included those of cell-cycle regulation, mitotic division and translation [34]. Their analysis followed the change from baseline and grouped stable disease with response, whereas lisaR compared CR/PR with PD at each time point, so the correspondence concerns direction. The mitochondrial signal higher in PD during treatment is an observation of this analysis.

### An example of lisaR in proteomics

To show that the same categories serve other molecular layers, we applied lisaR to protein abundance in clear cell renal cell carcinoma. Clark et al. characterised tumours and adjacent non-tumour tissue of this cancer by proteogenomics within CPTAC [35]. We used the 80 patients with matched tumour and NAT samples and compared tumour with NAT within each patient, so that positive NES points towards the tumour (Fig. 4A).

**Figure 4.**
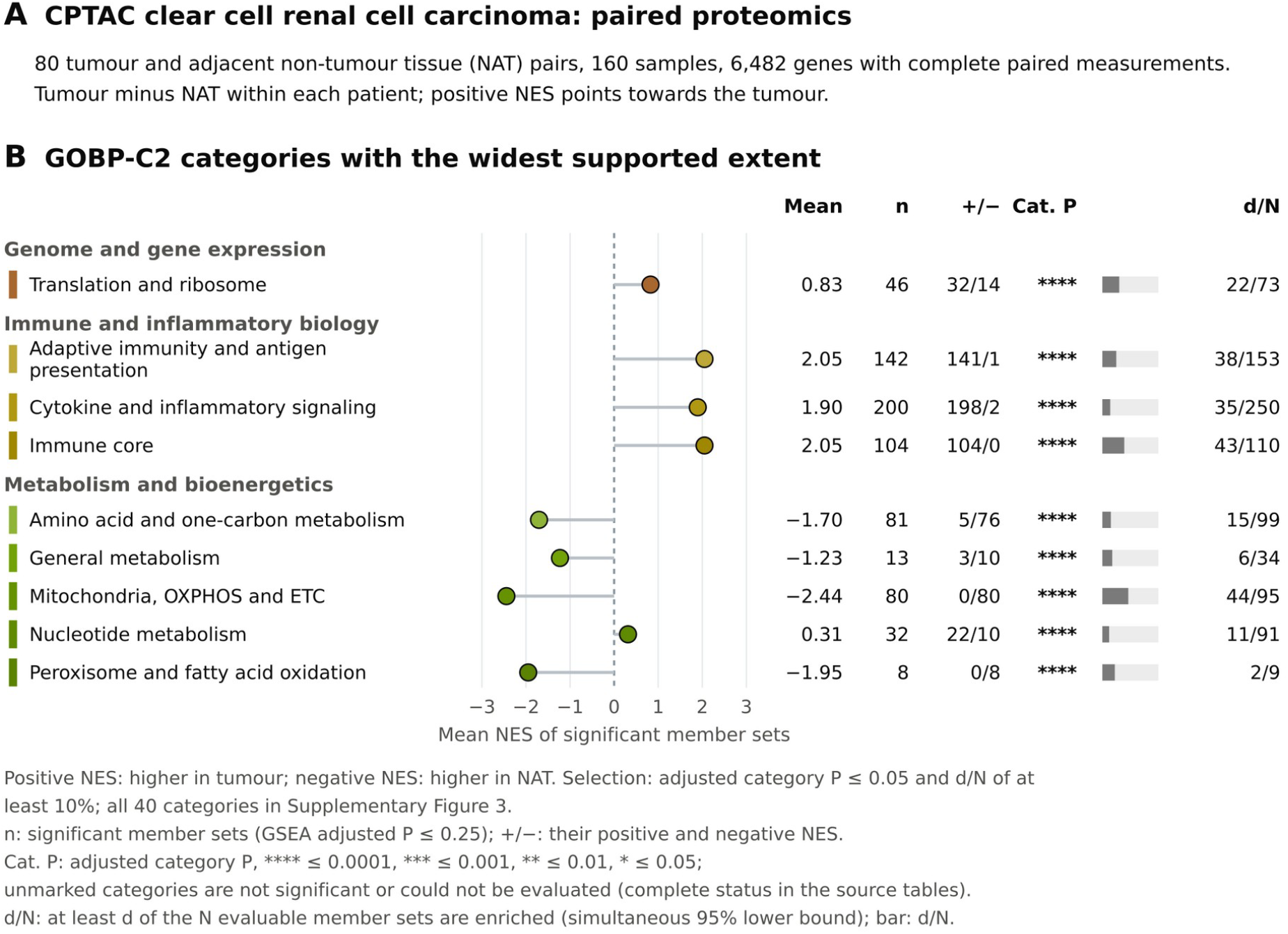
Category profile of paired tumour and adjacent-tissue proteomics in clear cell renal cell carcinoma. **A**. Design: 80 matched pairs of tumour and adjacent non-tumour tissue (NAT), 160 samples and 6,482 genes with complete paired measurements [35]. The comparison is tumour minus NAT within patients, and positive NES points towards the tumour. **B**. The nine GOBP-C2 categories selected with the rule of Figure 3B; columns as in Figure 3B. All 40 categories are shown in Supplementary Figure 3.

Here, 31 of the 40 GOBP-C2 categories showed evidence of enrichment (Supplementary Figure 3), and nine had lower bounds covering at least 10% of their evaluable sets (Fig. 4B). Mitochondria, OXPHOS and ETC had the widest supported extent: at least 44 of its 95 evaluable sets were enriched, and all 80 of its significant sets had negative NES, with a mean of −2.44, pointing towards NAT. Amino acid and one-carbon metabolism (at least 15 of 99; 76 of 81 significant sets negative), Peroxisome and fatty acid oxidation (2 of 9; all 8 negative) and General metabolism (6 of 34; 10 of 13 negative) pointed in the same direction.

The immune categories pointed towards the tumour, led by Immune core, with at least 43 of 110 sets enriched and all 104 significant sets positive (mean 2.05). In PATHWAYS, six categories showed evidence of enrichment, each with every evaluable member set pointing towards the tumour, including Type I interferon signaling (at least 2 of 5 evaluable sets) and T-cell receptor signaling (2 of 12).

Translation and ribosome shows why the direction is read from the member sets. At least 22 of its 73 evaluable sets were enriched, one of the widest extents in this analysis, but its 46 significant sets split into 32 positive and 14 negative, with a mean NES of 0.83. The inference establishes that many of its sets are enriched; the sign counts show that they point in both directions.

This profile agrees in direction with the original study, which found lower levels of TCA-cycle, fatty-acid-metabolism and OXPHOS proteins in tumours than in NAT and higher levels in immune-response and interferon pathways, among others [35]. Clark et al. compared the two tissues as unpaired groups, and our analysis is paired. The profile of Amino acid and one-carbon metabolism, which also pointed towards NAT, is an observation of this analysis.

## Discussion

lisaR organises enrichment results from a broad range of gene-set sources into biologically meaningful categories that stay the same from one analysis to the next. In the two examples, the same GOBP-C2 categories organised RNA-seq from melanoma biopsies and paired proteomics from renal tumours, and every category led back to its member sets and genes.

Because the dictionaries were built before any study was analysed, they could be evaluated once, and each analysis applies them on the local computer without a language model.

Four properties shape what lisaR offers. Its breadth lets one vocabulary read Gene Ontology and curated pathway collections together: the core dictionaries classify 13,511 gene sets, and in the examples 93% of the significant GOBP-C2 sets received a category. Its categories are fixed within a version, so a category means the same in every study, and results obtained at different times can be compared descriptively on the same terms, as the pretreatment and on-treatment melanoma profiles showed. Its category inference turns a list of significant categories into a statement of extent. In both examples most GOBP-C2 categories passed the adjusted-P threshold, and the simultaneous lower bound identified the immune categories on treatment in melanoma and Mitochondria, OXPHOS and ETC in renal tumours as the categories with the widest supported extent, while the mean NES and its sign counts gave the direction of the underlying sets. Because the bounds rest on closed testing, they can be read together across all the categories of an analysis without a further correction [25].

Finally, its report travels as a set of web pages that anyone can open in a browser, and each figure carries the data and the code that produced it.

This design sits among complementary strategies (Table 1). Similarity-based methods such as REVIGO, EnrichmentMap, simplifyEnrichment and vissE build groups from the terms present in each result, so the summary adapts to that result [4, 8, 7, 9]. Predefined vocabularies such as GO slims, Categorizer and thematicGO fix the grouping in advance [12, 13, 14]. lisaR follows this second approach and, compared with these vocabularies, extends it to the curated pathway collections of MSigDB, which do not share a common ontology with GO. Language-model tools such as SummArIzeR, MAPA and EnrichGT annotate the groups found in each analysis [16, 19, 20]; lisaR uses the models once, to build the vocabulary, so that the same annotation can be evaluated and versioned before use. GeneTonic and GeneFEAST share with lisaR the aim of keeping gene-level evidence close to the summary [11, 10], and mitch addresses a related statistical question, enrichment across several contrasts at once [15].

Two properties of the vocabulary matter when a result is read. Its resolution is fixed within a version: a broad category brings related sets together and can hide differences among them, which the member-set view restores. PATHWAYS classifies only the gene sets whose names identify one of its signalling systems, so it complements the broader GOBP-C2 reading. The vocabulary itself is versioned: changes in gene-set collections or in biological interpretation call for a new version, and lisaR records the version used in every run.

The same construction can be repeated to add categories or collections, and each new version can be evaluated in the same way before use. Users can also bring their own categories through custom dictionaries. Within a version, lisaR offers a stable way to read enrichment results, from categories down to the genes that support them.

### Software and resource versions

lisaR 1.0 (R package version 1.0.0) is distributed under GPL-3.0-or-later and archived at Zenodo (https://doi.org/10.5281/zenodo.23044930). The dictionaries are lisa_dictionary_core 1.0.0, lisa_dictionary_expanded 1.0.0 and lisa_category_map 1.0.0, and the human gene-set memberships are prepared from MSigDB 2026.1. Both examples used these dictionaries. The melanoma enrichment was run with R 4.5.3, fgsea 1.36.2 and lisaR 1.0, and the category inference of both examples with lisaR 1.0 and the hommel package 1.8. External gene-set memberships, pathway material and study data keep their own terms of use.

## Data availability

The melanoma RNA-seq data are available from GEO under accession GSE91061 [34], and the clear cell renal cell carcinoma proteomic data from the CPTAC Proteomic Data Commons under study PDC000127 [35]. The prepared example inputs used here are distributed in release v1.0.0 of https://github.com/DBM-OlmedaLab/lisaR-example-data as riaz-gse91061-lisa-examples-v1.0.0.tar.gz and cptac-ccrcc-lisa-prepared-v1.0.0.tar.gz; install_lisa_example_bundle() downloads them and verifies their checksums.

## Acknowledgements

We thank Dr María S. Soengas (Spanish National Cancer Research Centre, CNIO, Madrid, Spain) for critical reading of the manuscript and advice on the design of lisaR, and the other members of the Melanoma Group at CNIO and of the Drivers and Biomarkers of Metastasis Laboratory for helpful discussions. We thank Sergio Oterino-Sogo (Centre de Bioinformatique de Bordeaux, CBiB, University of Bordeaux, Bordeaux, France) for his independent evaluation of the LISA dictionaries. We are grateful to the patients who took part in the studies used as examples and to the investigators who made their data publicly available. Data used in this publication were generated by the National Cancer Institute Clinical Proteomic Tumor Analysis Consortium (CPTAC).

## Declarations

### Author contributions

C.V.-L.: Software, Validation, Visualization, Writing - review & editing. G.d.l.V.-B.: Validation, Writing - review & editing. D.O.: Conceptualization, Methodology, Software, Formal analysis, Visualization, Writing - original draft, Writing - review & editing, Supervision, Project administration, Funding acquisition. All authors read and approved the final manuscript.

### Use of AI tools

Language models are part of the method: four open-weight models scored gene sets during the construction of the LISA dictionaries, and four further models provided the concordance assessment (Methods). No language model is used when lisaR analyses a study. C.V.-L. and D.O. wrote the lisaR code and later used AI coding assistants to optimise it. The authors wrote and edited the full text and take full responsibility for its content. AI language models assisted with grammatical review. No image-generation tools were used.

