## Supplementary material for "lisaR: An LLM-Inferred Semantic Annotation of biological categories for gene set enrichment analysis": Suplemental data

This supplement gives the details needed to understand and reproduce the LISA dictionaries, their evaluation, the statistics of the category analysis and the two example studies.

### S1. The LISA dictionaries

This section describes what the dictionaries contain and where they are found. The core and expanded dictionaries and the category map are installed with lisaR in the folder extdata/dictionaries of the package, and a catalogue that describes every category, with up to three representative gene sets, is installed in extdata/semantic-catalog. Each assignment gives the collection, the gene-set identifier and name, its source, the category identifier and display name, and the tier.

#### S1.1. Collections, sources and tiers

Four collections organise the gene sets. GOBP-C2 contains the GO Biological Process gene sets and the canonical pathways of MSigDB C2, GOMF the GO Molecular Function gene sets and GOCC the GO Cellular Component gene sets. PATHWAYS reads the gene sets of GOBP-C2 whose names identify a signalling system, so a gene set can appear in both GOBP-C2 and PATHWAYS (Table S1).

**Table S1. The four LISA collections in the core dictionary.**

| Collection | Categories | Supercategories | Core assignments | Core gene sets |
|---|---|---|---|---|
| GOBP-C2 | 40 | 10 | 11,729 | 10,743 |
| GOMF | 37 | 9 | 1,860 | 1,768 |
| GOCC | 12 | 6 | 1,003 | 990 |
| PATHWAYS | 69 | 17 | 921 | 813 |
| All collections | 158 | 42 | 15,513 | 13,511 distinct |

The 13,511 distinct gene sets of the core dictionary come from GO Biological Process (6,937), GO Molecular Function (1,768), GO Cellular Component (990), Reactome (1,701), WikiPathways (830), KEGG (805), BioCarta (273), PID (188) and three smaller canonical pathway sources (19). The core tier holds the assignments with a final consensus score of 4, and the expanded tier those with a score of 3 or 4, so the expanded dictionary contains the core; it has 23,226 assignments of 14,230 distinct gene sets. A gene set can belong to several categories, and 1,826 gene sets of the core dictionary have more than one.

The package licence covers the LISA material. MSigDB, KEGG and BioCarta keep their own terms of use, and the resource table installed with the dictionaries lists the conditions that apply to each source.

#### S1.2. Categories

The 158 categories are listed by collection and supercategory in Section S8, with the number of gene sets assigned to each in the core dictionary. The package catalogue adds the written definition of every category.

### S2. How the dictionaries were built

This section describes how the scores behind each assignment were obtained and how the assignments were selected from them.

#### S2.1. Categories and model inputs

We wrote a name, a core scope and exclusion notes for every category before any scoring. For GOMF and GOCC, the models received these definitions together with the identifier, the name and example genes of each gene set. For PATHWAYS, a name-first prompt placed the gene-set name, the title of the source pathway, its database and its identifier before the canonical genes, and asked for one integer score from 0 to 4 for each category, returned as JSON.

#### S2.2. The four language models

Four open-weight models scored every collection (Table S2). Qwen3.5 and Gemma 4 act as the anchor models of the consensus rule, and Mistral Small 3.2 and Phi-4 as supporting models.

**Table S2. Language models used to build the dictionaries.**

| Model | Identifier | Developer | Run with |
|---|---|---|---|
| Qwen3.5 27B | Qwen/Qwen3.5-27B | Qwen team, Alibaba Cloud | vLLM |
| Gemma 4 31B | google/gemma-4-31B-it | Google | vLLM |
| Mistral Small 3.2 24B | mistral-small3.2:24b | Mistral AI | Ollama 0.12.9 |
| Phi-4 14B | phi4:14b | Microsoft | Ollama 0.12.9 |

#### S2.3. Scoring scales

The scales ask how well the gene set fits the meaning of the category.

- **GOMF.** 0, no relationship; 1, a weak or contextual relationship; 2, a supported secondary function; 3, a major function; 4, a primary or canonical function.

- **GOCC.** 0, 3 or 4, judging the identity of the cellular compartment rather than a molecular activity.
- **PATHWAYS.** 3, a strong relationship with the named pathway, such as a related module or a receptor-ligand axis; 4, definitional or core identity. A score of 3 or 4 requires the name and source of the gene set to identify the pathway, and canonical genes can only reinforce that reading. Broad process, disease and downstream response terms enter only when their names identify the signalling system.

#### S2.4. Consensus rule for GOBP-C2, GOMF and GOCC

The rule takes the first row of Table S3 that the scores of a gene set and category satisfy. Gene sets that reach a final consensus score of 3 in no category remain unclassified.

**Table S3. Consensus rule for the Gene Ontology collections.**

| Anchor scores (Qwen3.5 and Gemma 4) | Supporting scores (Mistral Small 3.2 and Phi-4) | Final consensus score | Tier |
|---|---|---|---|
| Both 4 | Any | 4 | Core and expanded |
| Both at least 3, one of them 4 | At least one of 3 or more | 4 | Core and expanded |
| Both at least 3 | Any | 3 | Expanded |
| Both at least 2, one of them 3 or more | At least one of 3 or more | 3 | Expanded |
| Any other combination | | Below 3 | Unassigned |

In GOBP-C2, a missing model score was replaced by the most frequent of the available scores for the same gene set and category, taking the lower value when two scores were equally frequent; 3,649 gene-set/category pairs had at least one score supplied in this way, and the supplied values are labelled as such. In GOMF and GOCC, a pair that lacks a required score is left unscored.

#### S2.5. Selection rule for PATHWAYS

PATHWAYS has its own branches (Table S4). The anchor pair can come in either order, and a score observed from a model is distinguished from one supplied for a missing response.

**Table S4. Selection branches for PATHWAYS.**

| Anchor scores | Supporting scores | Pairs | Tier |
|---|---|---|---|
| 4 and 4 | Any | 824 | Core and expanded |
| 4 and 3 | Mistral Small 3.2 and Phi-4 both observed at 4 | 97 | Core and expanded |
| 4 and 3 | Both at least 3, exactly one of them 4 | 100 | Expanded |
| 3 and 3 | Mistral Small 3.2 and Phi-4 both observed at 4 | 34 | Expanded |

The core tier thus holds 921 pairs and the expanded tier 1,055. A pair with one anchor at 4 and the other below 3 is unassigned. PATHWAYS scored 11,653 gene sets. For eight of them the Phi-4 responses were missing, and the missing scores were replaced by the median of the available model scores and labelled as such.

#### S2.6. Versions

The released dictionaries are identified by their version, 1.0.0, and distributed with the package. The MSigDB release read during construction was not recorded; lisaR prepares its human gene-set memberships from MSigDB 2026.1.

### S3. Evaluation of the dictionaries

This section describes how the evaluation set was built and gives the human and model judgements in detail.

#### S3.1. The evaluation set

The set was designed to reach every category with core assignments while keeping apart the gene sets used during development. We started from the 23,953 human MSigDB gene sets other than Hallmarks, of which 13,511 have a LISA assignment, and excluded the 1,388 gene sets used while developing the construction rules. Gene sets were drawn in strata: coverage of the categories, the multilabel frontier of gene sets assigned to several categories, a random sample of assigned gene sets, and unassigned gene sets for the negative controls. We then reduced the sample to the assigned gene sets needed to reach all the categories it represented: those that were the only representative of a category in the sample, and the smallest further group, found by an exact set cover, that reached the remaining categories. Each control pairs an unassigned gene set with a category taken in the order of the category map, independently of the content of the gene set. The rows were presented in random order.

The final set has 217 relations from 132 gene sets and 147 categories. Its 198 core assignments belong to 113 gene sets: 187 come from the coverage stratum, eight from the multilabel frontier and three from the random stratum. They come from GOBP-C2 (87), GOMF (32),

GOCC (11) and PATHWAYS (68) and cover 145 of the 158 categories; the other two of the 147 categories appear only in controls. The remaining 19 relations are the negative controls.

#### S3.2. Human evaluation

Three evaluators, working independently, judged all 217 relations. Each relation was classified as central and direct, secondary and plausible, unsupported or indeterminate, and central and secondary judgements counted as support. The three evaluators supported all 198 core assignments and rejected all 19 controls.

#### S3.3. Concordance with four further language models

Four language models that took no part in the construction judged the same 217 relations with the same four-level scale: Claude Sonnet 5 and Claude Opus 5 (Anthropic) and GPT-5.6 Terra and GPT-5.6 Sol (OpenAI). Each model ran twice, so every relation received eight judgements, and a relation counted as rejected when at least four of them were unsupported.

Of the 198 core assignments, 196 were supported in all eight judgements, 156 of them always as central. Of the 1,584 judgements of assignments, 1,451 were central, 126 secondary and 7 unsupported. Two assignments received mixed judgements (Table S5). The models that rejected the deneddylase assignment distinguished the removal of NEDD8 from the ligase activity named by the category. All 19 controls were rejected, 18 of them in all eight judgements. The exception, HP_NEOPLASM_OF_THE_EYE paired with RET signaling, received one supporting judgement, so that 151 of the 152 control judgements were unsupported. The model panel complements the human evaluation on the same relations.

**Table S5. The two assignments with mixed model judgements.** Support counts the supporting judgements out of eight. S, supported; U, unsupported, for runs 1 and 2 of each model.

| Gene set | Category | Support | Rejected | Judgements by model |
|---|---|---|---|---|
| GOMF_DENEDDYLASE_ ACTIVITY | Ubiquitin and SUMO ligase activity | 2 | Yes | Sonnet 5 U/U; Opus 5 S/S; Terra U/U; Sol U/U |
| GOMF_NUCLEOSOMAL_ DNA_BINDING | Structural constituent activity | 7 | No | Sonnet 5 S/U; Opus 5, Terra and Sol S/S |

### S4. Custom dictionaries

Researchers can read enrichment results through their own categories with the same workflow, statistics and report. This section describes what a custom dictionary contains and how it enters a run.

#### S4.1. The three tables

A custom dictionary is made of three tab-separated tables (Table S6). Each gene set of the assignments must be present in the memberships, and each category in the category map.

**Table S6. Tables of a custom dictionary.**

| Table | Columns | Content |
|---|---|---|
| Gene-set memberships (TERM2GENE) | gs_collection, gs_subcollection, gs_name, gs_exact_source, gene_symbol | One row for each gene of each gene set |
| Assignments | universe, gene_set_id, gene_set_name, source_id, category_id, category_display_name, tier | One row for each gene set and category |
| Category map | category_id, display_name, macrogroup_id, macrogroup_name, macrogroup_order, category_order_within_macrogroup | One row for each category, with its supercategory and display order |

#### S4.2. Registration and validation

install_lisa_resource() registers each table in the resource registry of lisaR with its checksum, version, species and approved origin, so that every run records exactly which resources it used. validate_lisa_custom_resources() then checks the three registered tables together:

- the tier of every assignment is custom;
- the universe of every assignment is GOBP-C2, GOMF, GOCC or PATHWAYS;
- every assignment appears once;
- the category identifiers and labels agree with the category map;
- every gene set of the assignments is present in the memberships;
- the registered schema, species and checksums match the tables.

The validator checks the structure and the relationships between the tables. The biological soundness of the categories remains the responsibility of whoever defines them.

#### S4.3. Use in a run

The registered tables are selected in the study configuration, through the keys dictionary_resource, category_map_resource and term2gene_resource, or passed to run_lisa() through its resources argument. validate_lisa_config() confirms that the configuration and its resources are ready before the analysis. From that point, the category summaries, the category inference and the report work as with the LISA dictionaries, and the run records the resources it used. The package vignette "Custom dictionaries and supercategories" follows the procedure in eight steps, with an example of 48 assignments in four categories that is installed with the package in extdata/custom-resources.

### S5. Statistical details

This section gives the definitions behind the category summaries, the category inference and the gene-card scores. We start with an example.

#### S5.1. A worked example

In the on-treatment melanoma analysis, Immune core has 185 member gene sets in the dictionary, and three counts describe it (Table S7).

**Table S7. The counts that describe a category, for Immune core in the on-treatment melanoma analysis.**

| Count | Definition | Immune core |
|---|---|---|
| Mapped members | Gene sets assigned to the category | 185 |
| N | Members that can be tested: eligible, with a finite raw GSEA P value | 134 |
| n | Members in the descriptive summary: finite NES and GSEA adjusted P ≤ 0.25 | 117 |
| d | Simultaneous 95% lower bound on the number of enriched members among the N | 52 |

The adjusted category P value, $2.1 \times 10^{-43}$, states that Immune core shows evidence of enrichment, and d/N = 52/134 states that at least 52 of its 134 testable members are enriched. The mean NES of the 117 significant members, 2.01, and their signs, 115 positive, give the direction. The two summaries use different members and answer different questions, and the inferential proportion is d/N. A category can also lack evidence. In the renal proteomics, EMT and cell-state plasticity has 34 mapped members and 11 testable ones, an adjusted category P of 1, d = 0 and no significant members. A bound of 0 indicates insufficient evidence of enrichment rather than its absence.

#### S5.2. Ranking and enrichment

The ranking statistic is signed and refers to the same comparison as the effect sizes, and every tested gene enters the ranking. Gene identifiers are matched to canonical symbols. A gene without a ranking value keeps a missing value, and duplicated symbols stop the analysis until their resolution is declared. lisaR runs multilevel GSEA with fgsea on the gene-set memberships of each selected collection [1]. The set-size limits apply to the genes that a set shares with the ranking, 10 to 600 by default, and the defaults also fix eps = 0 and a root random seed of 1729.

#### S5.3. Category summaries

For a category c, S(c) is the set of its distinct member gene sets with a finite NES and a GSEA adjusted P value at or below the cutoff, 0.25 by default. The category mean is the sum of their NES values divided by their number, n, and it is reported when n is at least 1. The positive and negative counts, and their percentages, use n as the denominator. The same-direction percentage is the positive percentage for a positive mean and the negative percentage for a negative mean. Where quartiles are shown, they use type-7 quantiles and describe the spread between member sets.

#### S5.4. Category inference

**The family.** For each differential expression analysis, the hypothesis family is the union of the eligible gene sets across GOBP-C2, GOMF, GOCC and PATHWAYS, including the sets without a LISA category and excluding Hallmarks (Table S8). The family is fixed before any filtering by significance or by the direction of the NES, and the test uses the raw GSEA P values. A null hypothesis tested in more than one collection, with the same gene set, membership, ranked genes and scoring context, enters once with the largest of its P values. Gene sets with different identifiers remain separate hypotheses even when they contain the same genes, so the number of hypotheses counts gene sets rather than independent mechanisms. An eligible gene set without a raw P value would enter with P = 1; every eligible gene set of the three analyses reported here had a raw P value.

**Table S8. Hypothesis families of the three example analyses.**

| Analysis | Hypotheses in the family | GOBP-C2 categories with adjusted P ≤ 0.05 |
|---|---:|---:|
| Melanoma before treatment, CR/PR minus PD | 10,360 | 24 of 40 |
| Melanoma on treatment, CR/PR minus PD | 10,361 | 30 of 40 |
| Renal proteomics, tumour minus NAT | 6,673 | 31 of 40 |

**The procedure.** lisaR passes the family to the hommel R package with its robust option, `hommel(p, simes = FALSE)` [2]. This applies closed testing with the robust local test of Hommel, which replaces the Simes inequality with the harmonic factor m·H_m, where H_m = 1 + 1/2 + … + 1/m, and controls the familywise error rate under any dependence between the gene sets, provided that each raw P value is valid [3, 4, 5]. For the N testable members of a category, the local test of their intersection gives the adjusted category P value, and a category with P ≤ 0.05 contains at least one enriched member. Closed testing already controls the error across all categories of the analysis, so these P values receive no further multiple-testing correction.

**The lower bound.** The same closed testing gives d, the largest number k for which the procedure rejects, at the 5% level, the hypothesis that fewer than k of the N members are enriched. d is therefore a one-sided 95% lower bound on the number of enriched members, and d/N its proportion. lisaR finds d by a binary search over these tests, using the same inclusive comparison with the 5% level as the category P. The bounds are simultaneous over all subsets of the family, so the bounds of all the categories of an analysis can be read together; each analysis is a separate family. Categories share gene sets, so their bounds cannot be added.

**Marks in the figures.** Stars give the strongest threshold that the adjusted category P reaches: **** ≤ 0.0001, *** ≤ 0.001, ** ≤ 0.01 and * ≤ 0.05. Unmarked categories are either not significant or not testable (N = 0), and the source tables give the status of every category. The stars encode only the category P; d/N, direction and effect size have their own columns.

For the two examples, the category inference was added with lisaR 1.0 to the saved GSEA results of each analysis, without repeating the enrichment.

#### S5.5. Comparison of two profiles

For each category, the comparison places the significant mean of each profile, A and B, and reports A minus B. When only one side has significant members, the other side can show a contextual value, the mean NES of its evaluable members before the significance cutoff, labelled as such. When neither side has significant members, the category is shown without values, and a difference is calculated only when both values exist; a missing mean stays missing. In the melanoma example, the formal comparison of the two time points is the time-by-response interaction of the upstream model (S6.1), and the profile comparison describes how the two category profiles differ.

#### S5.6. Gene-card scores

For each gene and category, let n be the number of distinct classified gene sets that contain the gene in their leading edge or over-representation overlap, m the mean of their available NES values and p their minimum available adjusted P value. Let l be the log2 fold change of the gene and q its DE adjusted P value. The two scores are:

```
a = abs(m)
b = -log10(max(p, 1e-300))
S = n * (1 + a) * b            # LISA support
e = abs(l)
f = -log10(max(q, 1e-300))
C = sqrt(max(S, 0)) * e * (1 + f)  # DE-weighted contribution
```

A missing m gives a = 0 and a missing p gives b = 1; a missing l gives e = 0 and a missing q gives f = 0. Both scores are dimensionless and use absolute values, and the card shows the signed gene effects alongside. The source gene sets are the classified sets passed to the gene-level step, without a GSEA adjusted-P cutoff; when over-representation analysis is run, its overlaps contribute to n and p.

### S6. Example studies

This section describes how the inputs of the two examples were prepared and how lisaR was configured.

#### S6.1. Melanoma cohort

The study is Riaz et al. (Cell, 2017), GEO accession GSE91061 [6]. The prepared subset contains patients with one biopsy before and one during treatment, a response of CR/PR or PD, and a known history of prior ipilimumab, and excludes Pt3, the PCA outlier reported in the original study. It has eight CR/PR and 18 PD patients; four of the eight CR/PR patients and nine of the 18 PD patients had progressed on prior ipilimumab.

The upstream DESeq2 analysis estimated size factors and dispersions and applied nbinomWaldTest with maxit = 10000 [7]. Entrez identifiers were kept during modelling, and symbols were added at export with AnnotationDbi and org.Hs.eg.db. When several rows mapped to the same symbol, the row with the highest baseMean was kept, and ties went to the lower Entrez identifier.

Each time point had its own model, ~ ipi_progressed + response_pd, whose negated response_pd coefficient gave the CR/PR minus PD comparison used by lisaR. A longitudinal model with patient fixed effects, ~ patient_id + on_indicator + on_by_ipi + on_by_pd, estimated the changes over time. Here on_indicator is 1 for on-treatment biopsies, response_pd is 1 for PD and ipi_progressed is 1 for patients who had progressed on ipilimumab; on_by_ipi and on_by_pd are the products of on_indicator with ipi_progressed and with response_pd. The response and prior-ipilimumab main effects are absorbed by the patient effects. Table S9 lists the contrasts.

**Table S9. Contrasts of the upstream DESeq2 models of the melanoma cohort.**

| Contrast | Coefficient weights | Positive direction |
|---|---|---|
| Change over time in CR/PR | on_indicator 1; on_by_ipi 0.5 | On treatment minus before treatment, in CR/PR |
| Change over time in PD | on_indicator 1; on_by_ipi 0.5; on_by_pd 1 | On treatment minus before treatment, in PD |
| Time-by-response interaction | on_by_pd −1 | Change in CR/PR minus change in PD |
| Response before treatment | response_pd −1, in the pretreatment model | CR/PR minus PD before treatment |
| Response on treatment | response_pd −1, in the on-treatment model | CR/PR minus PD during treatment |

The weights of 0.5 average over the two prior-ipilimumab groups, which are balanced within each response group. The lisaR report uses the two response comparisons and compares their category profiles, with A the on-treatment profile and B the pretreatment profile.

#### S6.2. Renal cell carcinoma proteomics

The study is Clark et al. (Cell, 2019), CPTAC study PDC000127 [8]. We used the processed protein abundance table of the original publication, whose normalisation and batch correction precede the model described here. The paired records identify 80 patients and 160 tumour and NAT aliquots, with both samples of each pair in the same TMT plex.

RefSeq features were mapped to genes in steps (Table S10). Each accession was first looked up with its version, and without the version only when the exact lookup failed, using AnnotationDbi 1.72.0 and org.Hs.eg.db 3.22.0. A retained feature had to resolve to one Entrez gene and one canonical symbol, with the source symbol either the canonical symbol or an unambiguous alias of the same gene. Genes represented by several features were excluded, and complete measurements in all 80 pairs were required.

**Table S10. Preparation of the proteomic input.**

| Stage | Features or genes |
|---|---:|
| Processed RefSeq features | 11,710 |
| Features mapped to one gene and symbol | 11,662 |
| Genes represented by one feature | 10,755 |
| Genes with complete measurements in all 80 pairs | 6,482 |

For each gene and patient, the reference intensity supplied with the table was subtracted from the tumour and NAT values, and the paired difference was formed. An intercept-only model was fitted with limma::lmFit and moderated with eBayes(trend = FALSE, robust = FALSE), using limma 3.66.0 under R 4.5.3 [9]. The coefficient is the mean paired tumour minus NAT difference, its moderated t statistic ranks the genes, and the gene-level P values were adjusted by the Benjamini-Hochberg method. The values entered the model as supplied, with no further transformation, imputation or batch correction.

#### S6.3. lisaR settings

Both examples used the assignments of the core dictionary 1.0.0 and the gene-set memberships of MSigDB 2026.1 [10, 11, 12, 13], with multilevel fgsea, set-size limits of 10 and 600 genes and a GSEA adjusted-P cutoff of 0.25 for the summaries. The on-treatment melanoma analysis ranked 21,520 symbols from 22,070 input rows by the DESeq2 statistic, with root seed 1729; the renal analysis ranked 6,482 genes by the limma moderated t. The renal run used the four LISA collections and one tumour minus NAT comparison, without Hallmarks, over-representation analysis or KEGG maps. Its STANDARD report shows up to 15 gene sets and 40 genes for each category, and the melanoma category-evidence pages up to 25 gene sets and 40 genes.

### S7. Other tools for organising enrichment results

This section summarises, from their publications, how ten of the tools that we cite organise and present enrichment results. Each answers its own need, and several combine strategies. The descriptions refer to the cited versions.

**simplifyEnrichment** (journal article, 2023) [14]. It clusters functional terms by a binary cut of a semantic or gene-overlap similarity matrix, starting from GO identifiers or from a term similarity matrix, and it can combine the results of several gene lists. Word clouds, with keyword enrichment, summarise each cluster, and the results are shown as similarity and multi-list heatmaps, with a Shiny interface for selecting terms.

**thematicGO** (bioRxiv preprint, version 1, 2026) [15]. It runs GO enrichment on a gene list and matches the names of the enriched terms to themes defined by curated keywords; a term can join several themes. A cumulative theme score sums the $-\log10$ P values of its terms. Theme and term plots, term-to-theme tables with the input genes and a theme network are available in Python and on the web.

**PAVER** (journal article, 2024) [16]. It groups pathways by hierarchical clustering of precomputed embeddings, starting from a table of pathway identifiers and enrichment metrics, and labels each cluster with the pathway closest to its mean embedding. Heatmaps and scatterplots compare several analyses, in R and on the web.

**mitch** (journal article, 2020) [17]. It tests rank-based enrichment across one or many contrasts, starting from differential expression tables or prescored gene-by-contrast data. A multivariate analysis of centred ranks, with Benjamini-Hochberg adjustment, orders the results by significance, effect size or variation across contrasts, and a self-contained interactive HTML report and PDF plots show the patterns across contrasts.

**GeneFEAST** (journal article, 2025) [18]. It starts from enrichment results, genes of interest and their measurements in one or several experiments, and builds term graphs from the shared genes of interest, divided into communities and meta-communities. Navigable HTML reports combine term-gene and experiment-gene heatmaps with links to gene and literature databases and exports, and optional user annotations can be added to the displays.

**EnrichGT** (journal article, 2026, version 2.2.0) [19]. It runs enrichment on gene lists or ranked data with Reactome, KEGG or custom gene sets and clusters the pathways of several databases by a pathway-by-gene matrix. A language model can add labels and summaries to the clusters, and the results are presented in interactive tables, word clouds and Quarto reports.

**SummArIzeR** (journal article, 2026) [20]. It runs enrichment through Enrichr or offline with custom sets, clusters the terms by walktrap on a network of shared genes, and labels the clusters through prompts to a language model; P values can be pooled across conditions. An interactive network, cluster tables with their terms and genes, and multi-condition heatmaps and bubble plots present the results.

**MAPA** (bioRxiv preprint, version 1, 2025) [21]. It embeds the names and descriptions of enriched pathways, clusters them into modules by Louvain or other methods, and asks a language model, supported by the literature, for the name, summary and confidence score of each module. R and Shiny views link modules, pathways and molecules, and reports are written in HTML, PDF, Word and Markdown.

**GeneTonic** (journal article, 2021) [22]. It combines a DESeqDataSet, its results, an enrichment table and gene annotation in a Shiny application, with gene-set networks and enrichment maps divided into communities and gene-set scores computed from gene-level effects. Gene and gene-set views, expression heatmaps, bookmarks exported to reports and the code of each plot support the interpretation.

**vissE** (journal article, 2024) [23]. It clusters gene sets into walktrap communities of an overlap network and summarises each community with word clouds built from the TF-IDF weighting of gene-set names. Gene-level statistics and protein-interaction networks can be displayed alongside the clusters.

Together, these tools serve complementary needs: shorter lists, maps of related gene sets, themes defined in advance, patterns across contrasts and generated interpretations. lisaR shares with several of them the report and the route to gene-level evidence, and its own contribution lies in a vocabulary fixed before the analysis across GO and curated pathways, and in the statistical test of each category.

### S8. The 158 LISA categories

Table S11 lists every category of the four collections, grouped by supercategory in the order of the category map, with the number of gene sets assigned to it in the core dictionary.

**Table S11. The 158 LISA categories.**

| Supercategory | Category | Core gene sets |
|---|---|---|
| **GOBP-C2** | | |
| Developmental and neural programs | Development and morphogenesis | 1,010 |
| | Neurogenesis and synapse | 791 |
| Genome and gene expression | DNA damage, repair and checkpoints | 295 |
| | RNA processing and RNP biology | 263 |
| | Transcription, chromatin and epigenetics | 416 |
| | Translation and ribosome | 116 |
| Immune and inflammatory biology | Adaptive immunity and antigen presentation | 328 |
| | Cytokine and inflammatory signaling | 535 |
| | Immune core | 185 |
| | Innate immunity, complement and PRR | 308 |
| Metabolism and bioenergetics | Amino acid and one-carbon metabolism | 251 |
| | Carbohydrate metabolism and glycolysis | 172 |
| | Detoxification and xenobiotic metabolism | 126 |
| | General metabolism | 51 |
| | Lipid and fatty acid metabolism | 637 |
| | Mitochondria, OXPHOS and ETC | 143 |
| | Nucleotide metabolism | 175 |
| | Peroxisome and fatty acid oxidation | 21 |
| Microenvironment, vascular and ECM | ECM and matrix remodeling | 183 |
| | Vascular, angiogenic and lymphatic biology | 151 |
| Proliferation and cell cycle | Cell cycle and proliferation | 555 |

| Supercategory | Category | Core gene sets |
|---|---|---|
| Signaling and cell communication | Core cell signaling | 714 |
| | Endocrine and hormonal pathways | 497 |
| | Growth factor and RTK pathways | 412 |
| | TGF-beta and SMAD signaling | 129 |
| | Wnt, Notch, Hedgehog and Hippo | 263 |
| Stress, proteostasis and cell death | Apoptosis and regulated cell death | 321 |
| | Autophagy, lysosome and phagosome maturation | 154 |
| | ER stress and unfolded protein response | 48 |
| | Heat shock and acute stress response | 17 |
| | Hypoxia and HIF programs | 31 |
| | Oxidative stress, redox and ROS | 116 |
| | Proteostasis, ubiquitin and proteasome | 214 |
| Structure, motility and plasticity | Adhesion, junctions and polarity | 260 |
| | Cytoskeleton and contractility | 419 |
| | EMT and cell-state plasticity | 34 |
| | Migration, invasion and motility programs | 215 |
| Transport, trafficking and homeostasis | Ion transport and homeostasis | 382 |
| | Transport and localization | 418 |
| | Vesicle, endocytosis, exocytosis and secretion | 373 |
| **GOMF** | | |
| Catalytic and metabolic enzyme functions | Amino-acid metabolic enzyme activity | 7 |
| | Antioxidant and redox activity | 54 |
| | Carbohydrate metabolic enzyme activity | 58 |
| | Detoxification and xenobiotic activity | 6 |
| | General catalytic activity | 134 |
| | General metabolic enzyme activity | 23 |
| | Lipid metabolic enzyme activity | 86 |

| Supercategory | Category | Core gene sets |
|---|---|---|
| Cell-cycle and developmental regulatory functions | Cell-cycle regulatory activity | 3 |
| | Developmental regulator activity | 1 |
| Gene-expression molecular functions | Epigenetic and chromatin activity | 73 |
| | RNA processing activity | 47 |
| | Transcription regulation activity | 41 |
| | Translation regulation activity | 9 |
| Generic or host-interaction functions | Binding activity | 525 |
| | Viral-host molecular function | 1 |
| Immune, ligand and receptor functions | Complement molecular function | 3 |
| | Cytokine, chemokine, growth-factor and hormone ligand activity | 12 |
| | Immune receptor and antigen-recognition activity | 38 |
| Proteostasis, stress and cell-death functions | Apoptosis and cell-death regulatory activity | 3 |
| | Protease and protease inhibitor activity | 40 |
| | Protein folding and degradation activity | 11 |
| | Stress-response molecular activity | 2 |
| | Ubiquitin and SUMO ligase activity | 28 |
| Signaling and regulatory functions | Enzyme regulator activity | 59 |
| | Intracellular signaling activity | 12 |
| | Kinase and phosphatase activity | 104 |
| | Receptor activity | 122 |
| | Scaffold and adaptor activity | 16 |
| Structure, adhesion and ECM functions | Adhesion molecular function | 12 |
| | Cytoskeleton and motor activity | 13 |
| | Extracellular matrix molecular function | 13 |
| | Structural constituent activity | 17 |

| Supercategory | Category | Core gene sets |
|---|---|---|
| Transport, vesicle traffic and sensing functions | Ion and physicochemical sensing activity | 4 |
| | Ion channel and pump activity | 86 |
| | Olfactory and taste receptor activity | 6 |
| | Transporter activity | 188 |
| | Vesicle trafficking activity | 3 |
| **GOCC** | | |
| Cell surface, adhesion and ECM compartments | Extracellular matrix compartment | 24 |
| | Junction, adhesion and cytoskeletal compartment | 209 |
| | Plasma membrane and cell-surface compartment | 115 |
| Immune and signaling complexes | Immune signaling and inflammatory complex compartment | 29 |
| Metabolic organelles | Mitochondrial and metabolic organelle compartment | 46 |
| Nuclear and gene-expression compartments | Nuclear and chromatin compartment | 137 |
| | Transcription, RNA and ribosome complexes | 140 |
| Proteostasis and generic complexes | Other generic protein/macromolecular complex | 42 |
| | Proteostasis complex compartment | 34 |
| Secretory and endomembrane compartments | Endolysosome, autophagy and intracellular vesicle compartment | 116 |
| | ER, Golgi and secretory-route compartment | 96 |
| | Extracellular vesicle and secretome compartment | 15 |
| **PATHWAYS** | | |
| RTK_RAS_MAPK | EGFR/ERBB signaling | 67 |
| | FGFR signaling | 42 |
| | VEGF/VEGFR signaling | 11 |

| Supercategory | Category | Core gene sets |
|---|---|---|
| | PDGF/PDGFR signaling | 15 |
| | MET/HGF signaling | 15 |
| | KIT signaling | 8 |
| | RET signaling | 6 |
| | Insulin/IGF signaling | 25 |
| | RAS activation | 16 |
| | RAF-MEK-ERK cascade | 43 |
| | ALK signaling | 7 |
| | NTRK/TRK signaling | 24 |
| STRESS_MAPK | p38 MAPK signaling | 12 |
| | JNK signaling | 9 |
| PI3K_AKT_MTOR | Class I PI3K signaling | 33 |
| | AKT signaling | 37 |
| | mTORC1 signaling | 11 |
| | mTORC2 signaling | 1 |
| | AMPK signaling | 6 |
| RHO_GTPASE_CYTOSKELETON | RAC1 signaling | 10 |
| | RHOA-ROCK signaling | 11 |
| | CDC42 signaling | 8 |
| | PAK signaling | 2 |
| | FAK/SRC adhesion signaling | 5 |
| | Actin regulatory signaling | 1 |
| TGF_BETA_SMAD | TGF-beta/SMAD signaling | 17 |
| | BMP/SMAD signaling | 8 |
| | Activin/Nodal signaling | 6 |
| WNT_NOTCH_HEDGEHOG | WNT/beta-catenin signaling | 37 |
| | Non-canonical WNT/PCP signaling | 6 |
| | NOTCH signaling | 34 |
| | Hedgehog signaling | 17 |
| HIPPO_MECHANOSIGNALING | Hippo/YAP/TAZ signaling | 7 |
| | Mechanotransduction signaling | 1 |
| CYTOKINE_JAK_STAT | Type I interferon signaling | 10 |

| Supercategory | Category | Core gene sets |
|---|---|---|
| | Type II interferon signaling | 6 |
| | IL6/JAK/STAT3 signaling | 9 |
| | IL2/STAT5 signaling | 11 |
| | IL10/STAT3 signaling | 5 |
| | IL12/IL23 signaling | 11 |
| INFLAMMATORY_NFKB | TNF/NF-kB signaling | 17 |
| | IL1/TLR-NF-kB signaling | 17 |
| | Canonical NF-kB signaling | 15 |
| | Non-canonical NF-kB signaling | 6 |
| INNATE_IMMUNE_SIGNALING | TLR signaling | 30 |
| | RIG-I/MDA5 signaling | 9 |
| | cGAS-STING signaling | 7 |
| | NLRP3 inflammasome signaling | 8 |
| GPCR_SECOND_MESSENGER | GPCR-cAMP-PKA signaling | 31 |
| | GPCR-PLC-calcium signaling | 16 |
| | PLC-gamma signaling | 2 |
| | Calcium/calcineurin/NFAT signaling | 12 |
| | S1P/EDG receptor signaling | 8 |
| T_CELL_B_CELL_RECEPTOR_SIGNALING | T-cell receptor signaling | 17 |
| | B-cell receptor signaling | 13 |
| | CD28 costimulation signaling | 7 |
| DEATH_RECEPTOR_SIGNALING | FAS/death receptor signaling | 10 |
| | TRAIL death receptor signaling | 6 |
| NUCLEAR_RECEPTOR_SIGNALING | Estrogen receptor signaling | 9 |
| | Androgen receptor signaling | 5 |
| | PPAR signaling | 8 |
| STRESS_RESPONSE_SIGNALING | NRF2/KEAP1 signaling | 11 |
| | PERK/ATF4 UPR signaling | 8 |
| | IRE1/XBP1 UPR signaling | 3 |
| | ATF6 UPR signaling | 4 |
| RTK_MATRIX_GUIDANCE_SIGNALING | Ephrin/EPH signaling | 11 |

| Supercategory | Category | Core gene sets |
|---|---|---|
| | DDR1/DDR2 collagen receptor signaling | 1 |
| | Semaphorin/Plexin/Neuropilin signaling | 8 |
| CHEMOKINE_GPCR_SIGNALING | CXCR4/CXCL12 signaling | 12 |

### S9. Supplementary figures

The three figures show all 40 GOBP-C2 categories of each analysis, grouped by supercategory in the order of the category map. Their columns are those of Figure 3B: the mean NES of the significant member sets (GSEA adjusted P ≤ 0.25), their number n, their positive and negative NES, the adjusted category P as stars and the simultaneous lower bound d/N, also drawn as a grey bar. Stars mark the significant categories only, and the source tables give the status of every category. Categories without significant member sets show no mean.

**Supplementary Figure 1.** Melanoma cohort on treatment (ON), CR/PR minus PD: all GOBP-C2 categories. Positive NES is higher in CR/PR.

**Supplementary Figure 2.** Melanoma cohort before treatment (PRE), CR/PR minus PD: all GOBP-C2 categories. Positive NES is higher in CR/PR.

**Supplementary Figure 3.** Clear cell renal cell carcinoma proteomics, tumour minus NAT: all GOBP-C2 categories. Positive NES is higher in tumour.

### Supplementary Figure 1

Riaz on treatment (ON), CR/PR minus PD: all GOBP-C2 categories

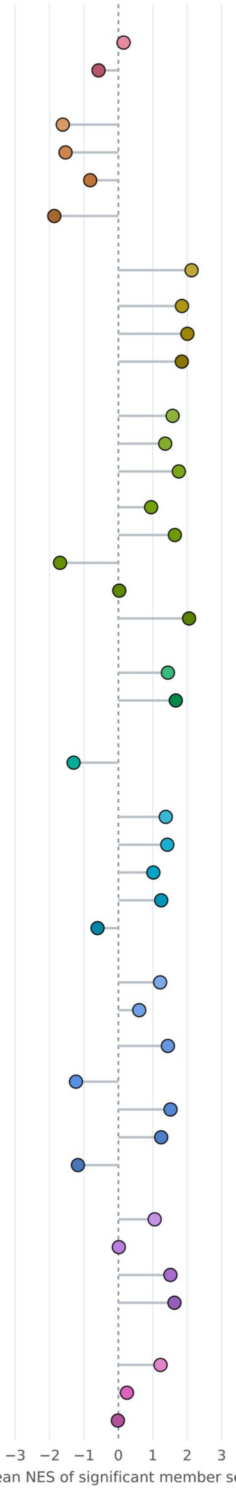


| Category | Mean | n | +/− | Cat. P | d/N |
|---|---|---|---|---|---|
| **Developmental and neural programs** | | | | | |
| Development and morphogenesis | 0.15 | 236 | 131/105 | **** | 8/716 |
| Neurogenesis and synapse | −0.58 | 144 | 43/101 | ** | 1/579 |
| **Genome and gene expression** | | | | | |
| DNA damage, repair and checkpoints | −1.62 | 146 | 5/141 | **** | 17/236 |
| RNA processing and RNP biology | −1.54 | 125 | 7/118 | **** | 12/185 |
| Transcription, chromatin and epigenetics | −0.82 | 146 | 39/107 | **** | 7/326 |
| Translation and ribosome | −1.86 | 60 | 0/60 | **** | 13/85 |
| **Immune and inflammatory biology** | | | | | |
| Adaptive immunity and antigen presentation | 2.13 | 238 | 237/1 | **** | 104/254 |
| Cytokine and inflammatory signaling | 1.85 | 376 | 372/4 | **** | 95/428 |
| Immune core | 2.01 | 117 | 115/2 | **** | 52/134 |
| Innate immunity, complement and PRR | 1.84 | 186 | 183/3 | **** | 38/223 |
| **Metabolism and bioenergetics** | | | | | |
| Amino acid and one-carbon metabolism | 1.58 | 92 | 87/5 | **** | 7/156 |
| Carbohydrate metabolism and glycolysis | 1.36 | 39 | 37/2 | * | 1/112 |
| Detoxification and xenobiotic metabolism | 1.76 | 64 | 64/0 | *** | 7/87 |
| General metabolism | 0.95 | 28 | 22/6 | **** | 4/41 |
| Lipid and fatty acid metabolism | 1.64 | 253 | 244/9 | **** | 31/460 |
| Mitochondria, OXPHOS and ETC | −1.70 | 67 | 4/63 | **** | 16/110 |
| Nucleotide metabolism | 0.03 | 61 | 32/29 | | 0/115 |
| Peroxisome and fatty acid oxidation | 2.06 | 5 | 5/0 | | 0/12 |
| **Microenvironment, vascular and ECM** | | | | | |
| ECM and matrix remodeling | 1.44 | 65 | 62/3 | *** | 2/142 |
| Vascular, angiogenic and lymphatic biology | 1.67 | 65 | 65/0 | **** | 6/107 |
| **Proliferation and cell cycle** | | | | | |
| Cell cycle and proliferation | −1.30 | 259 | 35/224 | **** | 47/417 |
| **Signaling and cell communication** | | | | | |
| Core cell signaling | 1.38 | 219 | 202/17 | **** | 21/555 |
| Endocrine and hormonal pathways | 1.42 | 142 | 135/7 | **** | 5/367 |
| Growth factor and RTK pathways | 1.02 | 68 | 56/12 | | 0/328 |
| TGF-beta and SMAD signaling | 1.25 | 28 | 26/2 | | 0/101 |
| Wnt, Notch, Hedgehog and Hippo | −0.61 | 54 | 16/38 | | 0/210 |
| **Stress, proteostasis and cell death** | | | | | |
| Apoptosis and regulated cell death | 1.21 | 90 | 77/13 | **** | 2/231 |
| Autophagy, lysosome and phagosome maturation | 0.61 | 26 | 17/9 | **** | 2/110 |
| ER stress and unfolded protein response | 1.44 | 3 | 3/0 | | 0/36 |
| Heat shock and acute stress response | −1.23 | 8 | 1/7 | | 0/14 |
| Hypoxia and HIF programs | 1.52 | 7 | 7/0 | | 0/21 |
| Oxidative stress, redox and ROS | 1.24 | 40 | 36/4 | * | 1/87 |
| Proteostasis, ubiquitin and proteasome | −1.17 | 72 | 8/64 | | 0/162 |
| **Structure, motility and plasticity** | | | | | |
| Adhesion, junctions and polarity | 1.06 | 103 | 82/21 | **** | 15/205 |
| Cytoskeleton and contractility | 0.01 | 162 | 85/77 | **** | 8/319 |
| EMT and cell-state plasticity | 1.51 | 4 | 4/0 | | 0/21 |
| Migration, invasion and motility programs | 1.63 | 92 | 86/6 | **** | 21/150 |
| **Transport, trafficking and homeostasis** | | | | | |
| Ion transport and homeostasis | 1.23 | 83 | 73/10 | **** | 6/272 |
| Transport and localization | 0.25 | 139 | 82/57 | **** | 7/293 |
| Vesicle, endocytosis, exocytosis and secretion | −0.01 | 87 | 43/44 | **** | 6/282 |

Positive NES: higher in CR/PR than in PD during nivolumab treatment.
n: significant member sets (GSEA adjusted P ≤ 0.25); +/−: their positive and negative NES.
Cat. P: adjusted category P, **** ≤ 0.0001, *** ≤ 0.001, ** ≤ 0.01, * ≤ 0.05;
unmarked categories are not significant or could not be evaluated (complete status in the source tables).
d/N: at least d of the N evaluable member sets are enriched (simultaneous 95% lower bound); bar: d/N.

### Supplementary Figure 2

Riaz pretreatment (PRE), CR/PR minus PD: all GOBP-C2 categories

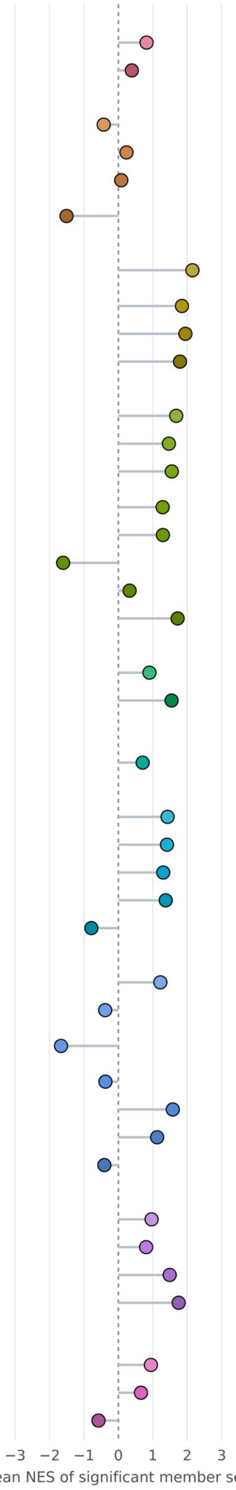


| | Mean | n | +/− | Cat. P | d/N |
|---|---|---|---|---|---|
| **Developmental and neural programs** | | | | | |
| Development and morphogenesis | 0.82 | 158 | 126/32 | **** | 9/717 |
| Neurogenesis and synapse | 0.39 | 61 | 39/22 | * | 1/580 |
| **Genome and gene expression** | | | | | |
| DNA damage, repair and checkpoints | −0.43 | 30 | 11/19 | | 0/235 |
| RNA processing and RNP biology | 0.24 | 32 | 18/14 | | 0/184 |
| Transcription, chromatin and epigenetics | 0.08 | 81 | 44/37 | | 0/324 |
| Translation and ribosome | −1.51 | 24 | 2/22 | *** | 5/85 |
| **Immune and inflammatory biology** | | | | | |
| Adaptive immunity and antigen presentation | 2.15 | 230 | 229/1 | **** | 96/254 |
| Cytokine and inflammatory signaling | 1.85 | 346 | 343/3 | **** | 57/427 |
| Immune core | 1.95 | 97 | 97/0 | **** | 39/136 |
| Innate immunity, complement and PRR | 1.79 | 140 | 137/3 | **** | 25/224 |
| **Metabolism and bioenergetics** | | | | | |
| Amino acid and one-carbon metabolism | 1.68 | 78 | 76/2 | **** | 5/156 |
| Carbohydrate metabolism and glycolysis | 1.47 | 31 | 30/1 | | 0/112 |
| Detoxification and xenobiotic metabolism | 1.55 | 36 | 35/1 | *** | 1/87 |
| General metabolism | 1.29 | 18 | 16/2 | **** | 3/41 |
| Lipid and fatty acid metabolism | 1.29 | 192 | 171/21 | **** | 6/459 |
| Mitochondria, OXPHOS and ETC | −1.61 | 53 | 4/49 | *** | 7/110 |
| Nucleotide metabolism | 0.33 | 20 | 12/8 | | 0/115 |
| Peroxisome and fatty acid oxidation | 1.72 | 5 | 5/0 | | 0/12 |
| **Microenvironment, vascular and ECM** | | | | | |
| ECM and matrix remodeling | 0.90 | 33 | 26/7 | | 0/142 |
| Vascular, angiogenic and lymphatic biology | 1.55 | 55 | 54/1 | ** | 4/108 |
| **Proliferation and cell cycle** | | | | | |
| Cell cycle and proliferation | 0.71 | 48 | 35/13 | **** | 1/418 |
| **Signaling and cell communication** | | | | | |
| Core cell signaling | 1.43 | 182 | 170/12 | **** | 19/556 |
| Endocrine and hormonal pathways | 1.41 | 95 | 90/5 | ** | 4/368 |
| Growth factor and RTK pathways | 1.30 | 50 | 46/4 | * | 1/327 |
| TGF-beta and SMAD signaling | 1.38 | 31 | 30/1 | | 0/101 |
| Wnt, Notch, Hedgehog and Hippo | −0.78 | 19 | 4/15 | | 0/210 |
| **Stress, proteostasis and cell death** | | | | | |
| Apoptosis and regulated cell death | 1.22 | 76 | 66/10 | *** | 2/232 |
| Autophagy, lysosome and phagosome maturation | −0.38 | 15 | 5/10 | *** | 2/110 |
| ER stress and unfolded protein response | −1.67 | 1 | 0/1 | | 0/36 |
| Heat shock and acute stress response | −0.38 | 5 | 2/3 | | 0/14 |
| Hypoxia and HIF programs | 1.58 | 8 | 8/0 | | 0/21 |
| Oxidative stress, redox and ROS | 1.13 | 20 | 17/3 | | 0/87 |
| Proteostasis, ubiquitin and proteasome | −0.41 | 11 | 4/7 | | 0/162 |
| **Structure, motility and plasticity** | | | | | |
| Adhesion, junctions and polarity | 0.97 | 75 | 58/17 | **** | 12/205 |
| Cytoskeleton and contractility | 0.81 | 70 | 56/14 | **** | 3/319 |
| EMT and cell-state plasticity | 1.49 | 2 | 2/0 | | 0/21 |
| Migration, invasion and motility programs | 1.75 | 69 | 69/0 | **** | 8/150 |
| **Transport, trafficking and homeostasis** | | | | | |
| Ion transport and homeostasis | 0.94 | 55 | 44/11 | ** | 3/272 |
| Transport and localization | 0.66 | 85 | 62/23 | | 0/292 |
| Vesicle, endocytosis, exocytosis and secretion | −0.58 | 67 | 21/46 | * | 1/282 |

Positive NES: higher in CR/PR than in PD before treatment.
n: significant member sets (GSEA adjusted P ≤ 0.25); +/−: their positive and negative NES.
Cat. P: adjusted category P, **** ≤ 0.0001, *** ≤ 0.001, ** ≤ 0.01, * ≤ 0.05;
unmarked categories are not significant or could not be evaluated (complete status in the source tables).
d/N: at least d of the N evaluable member sets are enriched (simultaneous 95% lower bound); bar: d/N.

### Supplementary Figure 3

CPTAC ccRCC, tumour minus NAT: all GOBP-C2 categories

| Category | Mean | n | +/− | Cat. P | d/N |
|---|---|---|---|---|---|
| **Developmental and neural programs** | | | | | |
| Development and morphogenesis | 0.83 | 96 | 73/23 | **** | 7/365 |
| Neurogenesis and synapse | 0.27 | 57 | 33/24 | **** | 4/272 |
| **Genome and gene expression** | | | | | |
| DNA damage, repair and checkpoints | 1.81 | 140 | 140/0 | **** | 9/162 |
| RNA processing and RNP biology | 1.36 | 96 | 86/10 | **** | 12/152 |
| Transcription, chromatin and epigenetics | 1.60 | 104 | 103/1 | **** | 6/213 |
| Translation and ribosome | 0.83 | 46 | 32/14 | **** | 22/73 |
| **Immune and inflammatory biology** | | | | | |
| Adaptive immunity and antigen presentation | 2.05 | 142 | 141/1 | **** | 38/153 |
| Cytokine and inflammatory signaling | 1.90 | 200 | 198/2 | **** | 35/250 |
| Immune core | 2.05 | 104 | 104/0 | **** | 43/110 |
| Innate immunity, complement and PRR | 1.83 | 103 | 103/0 | **** | 9/151 |
| **Metabolism and bioenergetics** | | | | | |
| Amino acid and one-carbon metabolism | −1.70 | 81 | 5/76 | **** | 15/99 |
| Carbohydrate metabolism and glycolysis | 0.22 | 40 | 22/18 | * | 1/86 |
| Detoxification and xenobiotic metabolism | −1.51 | 34 | 2/32 | * | 1/55 |
| General metabolism | −1.23 | 13 | 3/10 | **** | 6/34 |
| Lipid and fatty acid metabolism | −1.26 | 119 | 19/100 | **** | 17/291 |
| Mitochondria, OXPHOS and ETC | −2.44 | 80 | 0/80 | **** | 44/95 |
| Nucleotide metabolism | 0.31 | 32 | 22/10 | **** | 11/91 |
| Peroxisome and fatty acid oxidation | −1.95 | 8 | 0/8 | **** | 2/9 |
| **Microenvironment, vascular and ECM** | | | | | |
| ECM and matrix remodeling | 0.64 | 25 | 17/8 | | 0/79 |
| Vascular, angiogenic and lymphatic biology | 1.28 | 26 | 23/3 | * | 2/72 |
| **Proliferation and cell cycle** | | | | | |
| Cell cycle and proliferation | 1.74 | 169 | 169/0 | **** | 21/257 |
| **Signaling and cell communication** | | | | | |
| Core cell signaling | 1.59 | 160 | 158/2 | **** | 4/378 |
| Endocrine and hormonal pathways | 0.45 | 64 | 41/23 | ** | 1/192 |
| Growth factor and RTK pathways | 1.59 | 129 | 128/1 | | 0/225 |
| TGF-beta and SMAD signaling | 1.54 | 8 | 8/0 | | 0/57 |
| Wnt, Notch, Hedgehog and Hippo | 1.58 | 26 | 26/0 | | 0/97 |
| **Stress, proteostasis and cell death** | | | | | |
| Apoptosis and regulated cell death | 1.68 | 106 | 104/2 | **** | 9/162 |
| Autophagy, lysosome and phagosome maturation | 0.38 | 24 | 15/9 | *** | 1/92 |
| ER stress and unfolded protein response | 1.62 | 8 | 8/0 | | 0/25 |
| Heat shock and acute stress response | 1.58 | 4 | 4/0 | | 0/12 |
| Hypoxia and HIF programs | 1.64 | 10 | 10/0 | | 0/14 |
| Oxidative stress, redox and ROS | 0.65 | 25 | 17/8 | | 0/74 |
| Proteostasis, ubiquitin and proteasome | 1.49 | 91 | 89/2 | **** | 3/142 |
| **Structure, motility and plasticity** | | | | | |
| Adhesion, junctions and polarity | 1.42 | 71 | 64/7 | **** | 12/150 |
| Cytoskeleton and contractility | 1.33 | 89 | 84/5 | **** | 6/243 |
| EMT and cell-state plasticity | NA | 0 | 0/0 | | 0/11 |
| Migration, invasion and motility programs | 1.50 | 49 | 46/3 | *** | 4/100 |
| **Transport, trafficking and homeostasis** | | | | | |
| Ion transport and homeostasis | −1.14 | 56 | 10/46 | **** | 6/133 |
| Transport and localization | −0.30 | 86 | 36/50 | **** | 6/199 |
| Vesicle, endocytosis, exocytosis and secretion | 0.36 | 59 | 37/22 | ** | 2/228 |

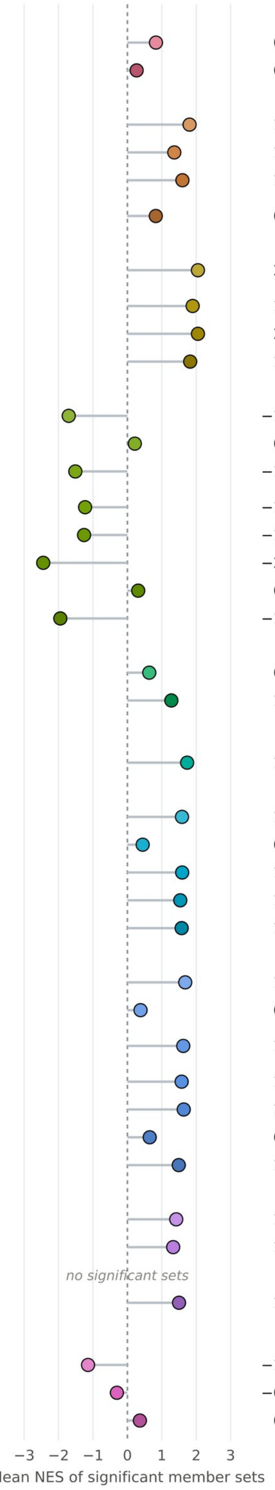


Positive NES: higher in tumour; negative NES: higher in NAT.
n: significant member sets (GSEA adjusted P ≤ 0.25); +/−: their positive and negative NES.
Cat. P: adjusted category P, **** ≤ 0.0001, *** ≤ 0.001, ** ≤ 0.01, * ≤ 0.05;
unmarked categories are not significant or could not be evaluated (complete status in the source tables).
d/N: at least d of the N evaluable member sets are enriched (simultaneous 95% lower bound); bar: d/N.
